# Spectral Separation of Evoked and Background Cortical Activity – Application to Short Latency Median Nerve Evoked Potentials

**DOI:** 10.1101/2021.09.09.459566

**Authors:** G. Fischer, D. Baumgarten, M. Kofler

## Abstract

**Background:** Spectral analysis of repeatedly evoked potentials (EPs) is challenging since recordings contain a superposition of evoked signals and background activity. We developed a novel approach, *N*-interval Fourier Transform Analysis (*N*-FTA), which allows for reliable separation and accurate assessment of evoked and background spectral components. We applied this approach to spectral analysis of median nerve short latency EPs for identifying spectral bands of clinical relevance.

**Methods:** We performed right median nerve stimulation in two volunteers at 2.46 Hz and 3.95 Hz stimulation rate (600 and 1000 repetitions respectively). We applied *N*-FTA for splitting the periodically repeated evoked components from irregular background activity and investigated spectral components in the low, medium and high frequency (LF-, MF-, HF-) bands. We present a signal processing approach, which allows for accurately extracting diagnostically relevant features (signal morphology, latency and amplitude) using an 18 Hz to 240 Hz bandwidth.

**Results:** In the LF-band *<* 10 Hz, evoked-to-background ratio (EBR) was below −15 dB due to the high level near DC background activity (1/f-activity, eye blinks, *θ*-band). Highest EBR was near −5 dB and obtained at a few tens of Hz. Relatively broad continuous segments of evoked activity were detected in the MF band. These frequencies of a few hundred Hz were linked to the signal segment between the *N*20 and *P*25 peaks in the time domain. High frequency oscillations (HFOs) near 600 Hz were approximately −25 dB below the background level and near the limit of the sensitivity of *N*-FTA. By subtracting stimulation artifacts and applying zero-phase filters it was possible to extract diagnostically relevant short latency EP features (signal morphology, latency and amplitude) only from the MF-band with a similar accuracy as a routinely used broad-band setting. HFOs displayed amplitudes of a few tenths of µV.

**Summary:** *N*-FTA allows for accurate, simultaneous spectral analysis of evoked and background activity in individual trials. The approach allowed for identifying target frequency bands of high evoked activity and for tailoring signal processing such, that morphology and latency can be obtained at a significantly reduced bandwidth. This should allow for development of more robust and faster recording techniques in the near future.

## 1. Introduction

Short latency somatosensory evoked potentials (SEPs) were introduced into routine diagnosis more than four decades ago. However, despite clear theoretical advantages, e.g., for the investigation of neuronal conduction in multiple scleroses [1], their application is restricted since recordings are time consuming and require highly qualified personnel for obtaining reliable results. Thus, technical developments which take advantage of recent improvements in state-of-the-art neuro-potential amplifier technology [2] appear highly appreciated.

To date, routine SEP recordings are based on IFCN standards, which were established in the early 1990’s [3]. These standards are based on experience obtained from hardware which, due to the technology at that time, essentially used analog filters. However, analog filters always involve time shifts in the signals. Due to the diagnostic importance of accurate assessment of latency, a broad filter bandwidth (up to 3 kHz) was chosen for reducing time shifts to a few tenths of ms, but this approach renders the technology prone to various types of interference.

In the last decade, high frequency oscillations (HFO) obtained increasing scientific interest and diagnostic application [4, 5, 6]. However, their frequency content is typically in the range of 400 Hz to 800 Hz and, thus, within the bandwidth of the IFCN recommendation. “Classical” SEPs reflect evoked activity defined by polarity and latency and they contain significant frequency content below the HF-band [7]. Early MEG studies on tibial and sural nerve SEPs obtained the typical signal morphology at only 100 Hz band-width, but low pass filtering significantly affected latency [8]. Recent scientific studies using SEPs for source reconstruction [4, 9] limit their bandwidth to a few hundred Hz. A clinical study investigating tibial nerve SEPs in a large cohort of HSP patients reduced bandwidth to 500 Hz for easing recording [10].

Recently, we provided theoretical estimations for signal bandwidth of compound action potentials generated by longitudinal tracts [11]. We found bandwidth estimates of some hundred Hz. The major effect limiting the bandwidth of neuro-potentials was the temporal dispersion of a volley (i.e., the distribution in timing of activation as caused by fast and slow conducting fibers within a tract). Experimental assessment of signal band-width is challenging, since recordings contain a superposition of the evoked components, biological background activity and technical noise sources. An early study investigated the low frequency spectrum of slow evoked responses [12]. More recent work focused of intra-cortical HF-activity [2, 13]. Detailed knowledge on the frequency content of short latency SEPs is scare.

In this study, we present a novel computational method, which allows for reliable separation of evoked and background activity in the frequency domain and for accurate assessment of both spectral components. We hypothesize, that this separation will provide important insight (to various types of evoked signals), which is required for developing faster and more stable recording techniques in the future. In a first step we investigate accuracy by means of synthetic test data. We then focus our work on experimental data, investigating cortical short latency responses to median nerve stimulation. This type of signal was considered due to its significance in routine diagnostics and due to the ongoing research on various frequency bands of these signals [7, 6]. In particular, we aim to show that the medium frequency (MF) band (10 Hz to 300 Hz), contains the relevant diagnostic information for routinely investigating SEP latency and morphology. Furthermore, we present a signal processing technique which accurately extracts median nerve cortical SEPs from a restricted band in the MF range only.

## 2. Methods

### 2.1. Fourier Transform of Near Periodic Signals

The mathematical theory of response averaging has been established in the 1970s and 1980s. A researcher assumes that repetitive responses to a stimulus are of the same or similar morphology and extracts the common mean morphology of all repetitions. Rompelman and Ros provided a comprehensive two-part tutorial review [14, 15]. In Appendix A we provide a summary of their work, and illustrate findings by computer simulations. In their Tutorial part 1 [14] they assumed an identical shape of all *N* responses. By applying a Discrete Fourier Transform (DFT) to an interval of *N* identical repetitions a periodic signal was obtained. The chosen duration of the time window was *exactly* the multiple *N* of the evoked cycle *T*_*E*_ (i.e., the finite periodic interval, Appendix A.1).

We refer to a DFT performed within the finite periodic interval *NT*_*E*_, by the term *N*-interval Fourier Transform (*N*-FT). For a periodic evoked signal, all non-zero *N*-FT components are at the harmonics *kf*_*E*_ of the stimulation rate (or evoked frequency) *f*_*E*_. In contrast, irregular background activity occurs at all frequencies. Thus, background activity becomes accessible from the spectral components between the harmonics. At the evoked harmonics, stimulus associated and background activity are superimposed. In section 2.2 we present an *N*-interval Fourier Transform *Analysis* (*N*-FTA), which allows for splitting of both spectral components.

In repetitive real world recordings, both random and systematic variations occur (Tutorial part 2 [15]). In Appendix A.4 we analyzed the effect of such variations on *N*-FTA. Small variations in activation timing influenced high frequency harmonics (i.e. harmonics near and above 1 kHz) defining the major restriction. Thus, we limited our analysis to frequencies below 750 Hz.

### 2.2. N-Fourier Transform Analysis

The spectral analysis which we applied for splitting evoked and spectral components in the frequency domain involves two steps:

- Firstly, a standard DFT was performed by selecting a data segment containing *exactly N* repetitions. Hence, a complex amplitude spectrum *a*_*m*_(*f*_*m*_) was obtained. All data points of this complex *N*-FT contained background activity. Evoked activity was superimposed at the spectral harmonics (see also Appendix A.1).
- Secondly, evoked and background components were split by spectral analysis, as described below and in Appendix A.1. This analysis computed the mean spectral background levels within each evoked harmonic interval and performed a reliable detection of (sufficiently large) evoked components. Here, we requested a probability of < 1/500 for the false positive acceptance of evoked components.

We converted the amplitude spectrum to spectral power density (PSD) *w*_*m*_(*f*_*m*_)

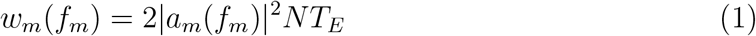

since, PSD allows for averaging power within intervals. The inverse of *NT*_*E*_ defines the frequency resolution in the spectrum. Furthermore, the phase angle *ϑ*_*m*_(*f*_*m*_) was obtained from the complex amplitude spectrum *a*_*m*_. Standard DFT software defines the time step zero at the first sample, while the observations from Appendix A.1 recommend selecting a representative morphological marker for time-point zero. Hence, we adjusted phase by a temporal offset *T* using the time shift theorem

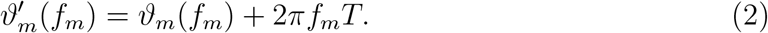

We set *T* to 20 ms (i.e. time zero near the N20 peak). Figure 1 illustrates the concept of averaging PSD within harmonic intervals. For obtaining the mean power of evoked activity components *e*_*k*_ and background components *b*_*k*_ the algorithm performs the following steps:

**Figure 1:**
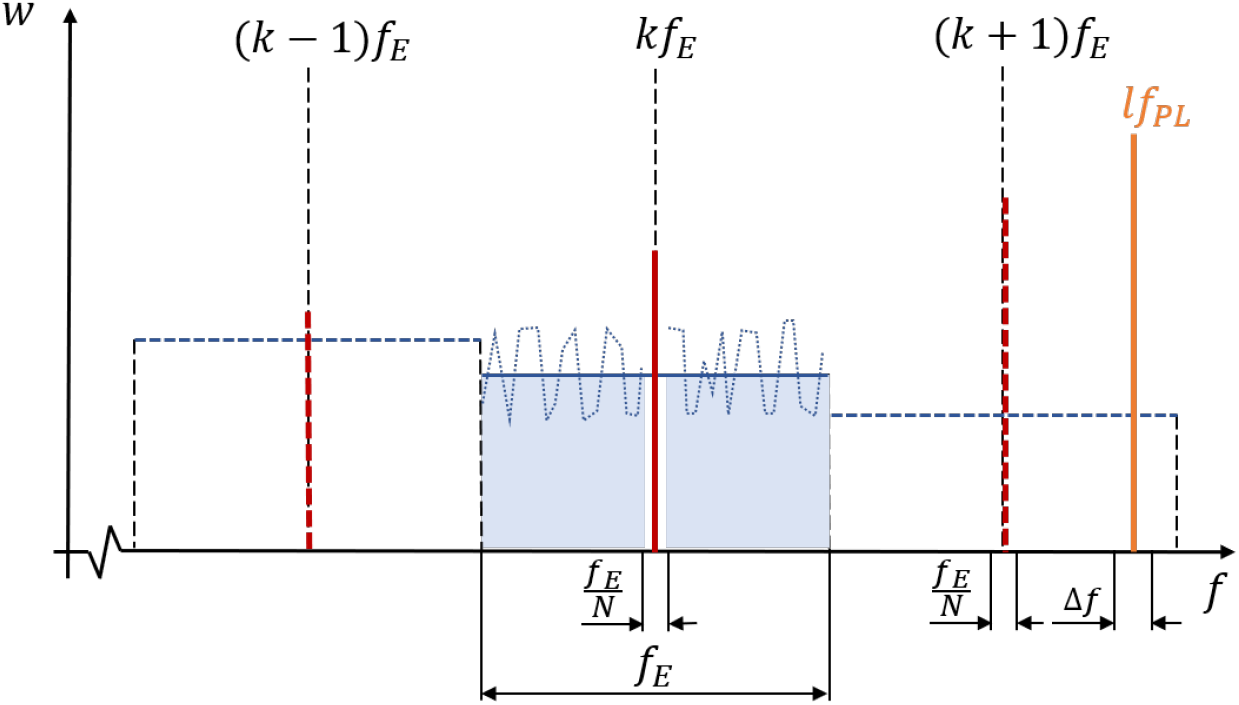
Illustration of the frequency intervals used for splitting of evoked and background activity (see text).

- Computation of *b*_*k*_ as the average background PSD within each evoked interval located at 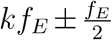. As can be taken from Figure 1, background components are all PSDs within an evoked interval except for the single value at the evoked harmonic *kf*_*E*_. In some intervals power line interference may occur. All harmonics (index *l*) of the power line frequency *f*_*PL*_ must be considered. For all power line harmonics *lP*_*PL*_ the adjacent intervals 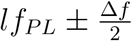 must be excluded from the analysis. We selected Δ*f*_*PL*_ = 0.5 Hz.
- For comparing PSD at the evoked harmonic *kf*_*E*_ with the background PSDs, two thresholds *p*_*L*_ and *p*_*H*_ were defined. We selected the 80% percentile of interval background PSDs as the lower threshold *p*_*L*_ and the 100% percentile (i.e. the highest background PSD in the interval) for the higher threshold *p*_*H*_.
- The spectral samples at the evoked harmonics *kf*_*E*_ contain a superposition of background activity and evoked activity. Here, we aim to identify harmonics in which a dominant contribution of the evoked component is statistically highly probable. Thus, we apply a two-step approach. In a first step, we select those samples at the harmonics with an amplitude above the threshold *p*_*L*_ as “candidates” *k*′ for evoked components. In a second step we apply the two amplitude/phase selection criteria (APSC, described below) for the decision on acceptance of a candidate. Thus, we obtain a set *K*_*A*_ of accepted harmonics.
- The PSD of the accepted harmonic samples *w*(*kf*_*E*_), *ϑ*′(*kf*_*E*_), *k* ∈ *K*_*A*_ reflects the power of *N* repetitions in the evoked responses. For obtaining a representation which is independent from the number of repetitions in an individual trial, we normalize evoked PSD by the number of repetitions

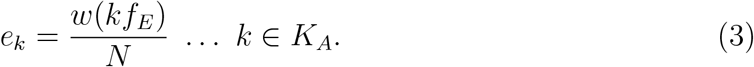

The computed power spectra *e*_*K*_ represent the PSD of a single averaged response.

#### Amplitude/Phase Selection Criteria (APSC)

- *APSC 1:* A candidate harmonic *k*′ is always accepted if *w*_*m*_(*k*′*f*_*E*_) *> p*_*H*_. In other words, if the amplitude of the candidate harmonic is larger than all background amplitudes in the interval, the evoked activity is likely to dominate background activity at that evoked harmonic. If we would average hypothetical white noise containing *N* segments, the probability for obtaining randomly the largest amplitude at a harmonic was 1*/N*. Thus, for 600 or 1000 repetitions in our trials the probability of accepting a false positive candidate was below the chosen 1/500 threshold.
- *APSC 2:* Furthermore, we investigate sets of three neighboring candidates {*k*′ −1, *k′, k′* + 1} which all three exceed the lower threshold *p*_*L*_ in amplitude. For a hypothetical white noise signal the probability to randomly detect such a set equals 1/5^3^ at the chosen 80% *p*_*L*_ level. For further reducing the probability of accepting false positive harmonics, we consider also phase information. It was observed from the example in Appendix A.1, that phase information of a repeated evoked response was reflected by a smooth function. For such a smooth function the phase 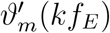 at the middle sample *k*′ should be approximately the mean 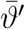 of its neighbor phases at *k*′ − 1 and *k*′ + 1. Thus, we request that the middle phase 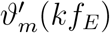 must not deviate by more than ±30° from 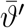 for acceptance of the set. Note that for a hypothetical white noise signal the probability to find a phase value within a 60° interval out of 360° equals 1/6. Thus, the probability for accepting a false positive set is *<* 1/5^3^ × 1/6 = 1/750.

Investigating a frequency interval below 750 Hz yields 304 harmonics at 2.46 Hz stimulation rate and 189 harmonics at 3.96 Hz stimulation rate. Thus, the expected value for false positive acceptances is statistically well below one for both acceptance criteria. Appendix A.2 provides a pseudocode listing of *N*-FTA.

#### Evoked-to-Background Ratio

We computed the evoked-to-background ratio (EBR) in dB as a frequency dependent signal-to-noise ratio

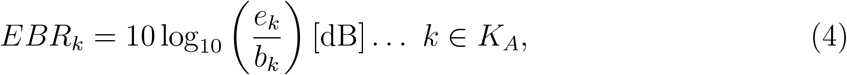

at the accepted harmonics *K*_*A*_.

### 2.3. Synthetic Test Data

For generating synthetic test data, a simulated compound action potential (CAP) was used. The data was taken from a previous study [11]. The implementation is described in detail in Appendices A.3 and A.4. Briefly, a reference CAP (peak-to-peak amplitude 0.32 µV, −6 dB bandwidth 50 Hz to 227 Hz) was obtained with a time step of 5 µs. Similar as in the experimental setting (see 2.4.1), *N* = 1000 repetitions were computed and three types of variations of individual responses were simulated: activation jitter (50 µs standard deviation), activation drift (0.5 ms over the entire time interval) and variation of amplitude (an initial amplitude scaling factor of 1.5 was linearly reduced to 0.5 after *N* repetitions). For re-sampling the data at 9.6 kHz we performed linear interpolation between the simulated time steps (5 µs) for accurately considering activation time shifts. Here, we investigated test data which contains all three types of variations simultaneously, similar as it is expected to occur in real world recordings. In Appendix A.4 each type of variation was further investigated independently. The simulated signal was also corrupted by white noise (standard deviation 0.1 µV).

### 2.4. Cortical Median Nerve SEPs

#### 2.4.1. Experimental Protocol and Equipment

The experimental protocol was approved by our institutional *Research Committee for Scientific Ethical Questions* and volunteers gave written informed consent. Due to the focus on algorithm development the number of volunteers was limited to two subjects A and B, both males of age 28 years, size 185 cm and 187 cm and body weight 72 kg and 78 kg, without any known neurological disease.

Two stimulation rates *f*_*E*_ were chosen for median nerve stimulation based on clinical and technical considerations: 2.46 Hz and 3.95 Hz, both avoiding interference with the power-line frequency (50 Hz) and its second harmonic (100 Hz). We first applied at least 600 stimuli for the lower stimulation rate (2.46 Hz) and then repeated stimulation, collecting at least 1000 stimuli for the higher stimulation rate (3.95 Hz). Subjects were asked to avoid movements and to remain physically relaxed during the recordings. We did not exclude any data from the segments (approximately 250 s each, corresponding to a frequency resolution of 4 mHz). Thus, the duration of a run was comparable to clinical routine recordings.

Sensory and motor thresholds were determined for monophasic (cathode) pulses of 200 µs duration. Stimulation was performed slightly above motor threshold, but below five times the sensory threshold. For the scope of methodological development, it was sufficient to perform only right median nerve stimulation. For generating different conditions of background activity, we asked subjects to open their eyes and blink naturally during the first trial, and to close eyes during the second trial.

For assessing background activity independently from stimulation, a single blinded trial was performed in each subject. Despite that no real stimulation, the subjects were told, that a sub-threshold stimulation was performed, for keeping their attention at a comparable level with the other trials. At the beginning of a blinded trial, the subject was asked to keep eyes open. After two minutes, they were asked to close their eyes. Two segments of 100 s duration each were used for studying cortical background activity in the absence of stimulation.

A DS7A neuro-stimulator (Digitimer Ltd., UK) was used for median nerve stimulation via a 30 mm pair of stainless steel electrodes (Spes Medica S.r.l., Italy). A digital function generator (AFG-2225, Good Will Instrument Co. Ltd, Taiwan) was used for triggering stimulation at selectable constant frequency. Signals were digitized at 9600 Hz using a g.USBamp (g.tec GmbH, Austria) biopotential amplifier. Gold plated Genuine Grass cup-electrodes (Natus Neurology, Ireland) were placed at the scalp positions Cz’, C3’, C4’ and Fz according to the international 10-20 system of electrode placement.

#### 2.4.2. Data Preprocessing

Figure 2 provides an overview of signal generation and processing steps. As for any SEP recording, repetitive stimulation evoked a series of responses while several interference sources corrupt the evoked target signal (derivation C3’-Fz for median nerve stimulation). Some of them are of biological origin, such as cortical background activity or eye blinks. Others are of technical origin, such as stimulation artifacts or power-line interference. Another sort of interference originates from the electrode-to-skin interface and is reflected by polarization potentials and ohmic noise. Importantly, all these types of interference carry individual spectral or temporal properties.

**Figure 2:**
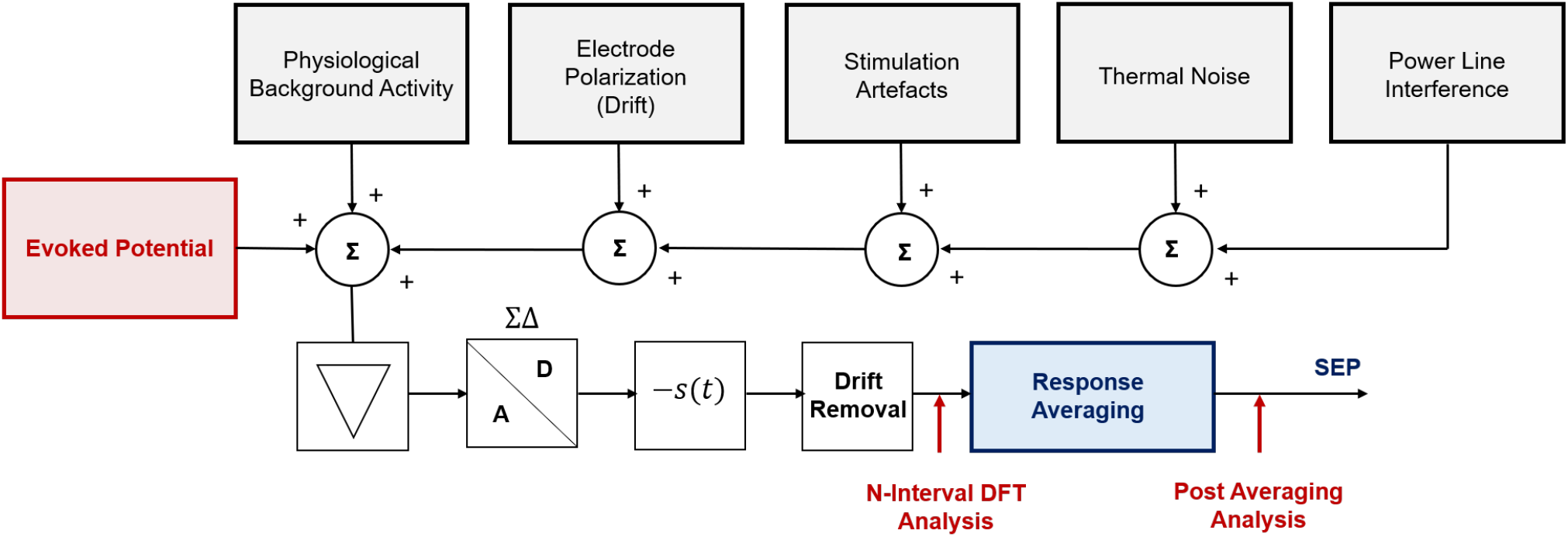
Overview on the spectral analysis concept applied in this study. We consider the evoked target signal (red) and a plurality of independent interference sources (gray). Each source of interference is characterized by different spectral properties. They are superimposed in the volume conductor and digitized by a state-of-the-art biopotential amplifier. Signal pre-processing removed stimulation artifacts in the time domain by subtracting a stimulus template *s*(*t*) and eliminated drift in the signals. *N*-FTA analysis was performed prior to response averaging.

##### Stimulation Artifact Cancellation

Stimulation artifacts are narrow pulses of high amplitude which repeat at the stimulation frequency *f*_*E*_. Thus, their frequency spectrum contains exactly the same evoked harmonics as the target signal. State of the art biopotential amplifiers provide a high resolution of amplitude in a broad range and accurately capture frequency components from “true DC” to the Nyquist-frequency. Thus, methods became available for removing stimulation artifacts by subtracting an artifact template from the signal [2]. Our approach for artifact subtraction is described in more detail in Appendix B. Briefly, an artifact template was generated by averaging all stimuli in a final pulse phase (approximately 1 ms to 5 ms after the stimulus). This stimulus template was subtracted from each stimulus. In the time interval between the last time step ahead of the stimulus and the final pulse phase, linear interpolation was applied.

##### Drift Cancellation

Varying electrode-to-skin contact causes drifting polarization voltages which are captured by true-DC biopotential amplifiers. DFT assumes a periodic repetition of the signal. Thus, a linearly drifting signal component is interpreted like a “saw-tooth” signal by a DFT which distorts the spectral profile. Therefore, linear drift was removed ahead of spectral analysis. Our algorithm estimated the drift in each trace by computing a linear interpolation from the first point to the last point in the finite periodic data segment (defined in section 2.1) and subtracted this linear function from the data.

#### 2.4.3. N-FTA Verification for Experimental Data

##### Evoked Component

As an alternative approach for spectral analysis of SEPs we computed a DFT of the averaged response and converted it to a PSD representation. Stimulation artifacts and linear drift were removed ahead of averaging. We did not apply any filtering, for not altering the frequency spectrum.

##### Background Component

As an alternative approach for assessing background activity we analyzed two 100 s segments from the single blinded trials. For investigating activity with open eyes we selected the 100 s interval starting 15 s after the begin of the recording. For investigating activity with closed eyes, we selected a 100 s interval terminating 15 s before of the end of the recording. For both segments we performed a DFT and computed the PSD. We averaged PSD in intervals centered at multiples of 2.46 Hz (analogous as in Figure 1). Furthermore, we excluded intervals adjacent to powerline harmonics *lf*_*PL*_ as described above. This allowed for direct comparison with background activity as obtained from *N*−FTA.

#### 2.4.4. Short Latency SEPs

##### Standard Setting

The standard setting applies comparable signal processing as for routine evaluation. Here, stimulation artifacts were **not** subtracted from the data. The drift removal component in Figure 2 was implemented as a 3 Hz to 2000 Hz 2^nd^-order Butterworth band-pass. Corner frequencies were chosen within the recommended range [3, 16]. Finally, SEPs were extracted by response averaging.

##### MF-Band Setting

The MF-band setting involved two steps for enhancing and assessing the quality of the evoked responses:

- Stimulation artifact cancellation was performed as described in section 2.4.2.
- The drift removal component in Figure 2 was implemented as a narrow zero-phase band-pass filter as described below.

We applied an 18 Hz to 240 Hz bandpass for extracting the most relevant signal components from the MF band. The selection of filter types was based on the following considerations. Since latency is the key diagnostic information in SEPs we used zero-phase filters for avoiding time shifts. For preserving signal morphology, we chose filter types with no or only moderate oscillations in their impulse response. For the low-pass component of the filter we used a 4^th^-order moving average filter. For the high-pass component we used a 2^nd^-order Butterworth filter with a corner frequency of 14.4 Hz. For obtaining a zero-phase characteristic we used the Matlab routine filtfilt which applies “forward and backward filtering”. The effective −3 dB corner frequency was at 18.0 Hz, due to applying the high-pass filter twice (see Appendix C.1).

##### Signal Quality Assessment

For verifying whether the MF band setting provides a comparable signal morphology and comparable latency to the standard setting (despite a significantly reduced bandwidth) we assessed the following parameters: for each trial the coefficient of correlation *r* was computed comparing the traces obtained from the MF band setting and the standard setting within the diagnostically relevant window (15 ms to 50 ms). Furthermore, we investigated the difference in latency between the two settings for the cortical SEP components N20 and P25.

For estimating the signal-to-noise ratio (SNR) of averaged responses, we assumed a relatively early appearance of the short latency response in each evoked cycle, while in the later phase of each cycle near-isoelectric segments may be expected, which may contain residual background activity or noise. We assumed that for the slower stimulation rate of 2.46 Hz the late time interval of 325 ms to 375 ms may be suited for noise floor estimation and we restricted analysis of the SNR to the slower stimulation rate. We assessed the root-mean-square noise amplitude *n*_*r*_ from this noise floor segment.

An early segment in the cycle reflected a superposition of the evoked target signal and residual noise. We chose a target signal interval of 15 ms to 30 ms. Within this target interval we assessed the signal level *σ*_*r*_ (rms amplitude of the superposition of the target signal and residual noise). We assumed independent (or in a mathematical sense orthogonal) morphology [17] of target signals and noise components. We then obtained an estimate for the rms-SNR

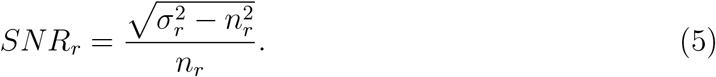

#### 2.4.5. High Frequency Oscillations

For obtaining HFOs, the drift removal block in Figure 2 was implemented by a zero-phase band-pass identically as described in [6]. Briefly, a 3^rd^ order Butterworth 400 Hz to 800 Hz band-pass filter kernel was obtained and filtering was applied once in a forward time direction and once in a backward time direction (Matlab routine filtfilt). This double application of the filter slightly narrows bandwidth yielding the actual −3 dB corners at 419 Hz and 765 Hz (see Appendix C.1). We compared the peak-to-peak amplitude of the averaged HFO-responses within the 10 ms to 30 ms interval (near N20 evoked oscillation) to the peak-to-peak amplitude within the 40 ms to 60 ms interval (background amplitude). For frequency domain analysis, we investigated the 450 Hz to 750 Hz band of evoked activity as obtained from *N*-FTA.

## 3. Results

### 3.1. Synthetic Test Data

A comparison of *N*-FTA evoked and background activity of a simulated CAP (see section 2.3) with reference values is shown in Figure 3. Evoked activity was accessible within a frequency band of 16 Hz to 363 Hz. The frequency resolution was defined by the evoked rate *f*_*E*_ = 3.95 Hz. For the *N* = 1000 stimuli, evoked spectral components, which were approximately a factor of 1000 or more below the background level, were not captured. The deviation between the evoked *N*-FTA components and the reference was −0.06 dB ±0.72 dB (mean ±standard deviation), with highest deviations at low signal levels. For the background activity we investigated the frequency band from the evoked frequency to 750 Hz and compared it to the background level corresponding to the white noise level (see Appendix A.3). The deviation was 0.09 dB ± 0.24 dB. At frequencies near the peak of the evoked components we observed a moderate increase of the background level by less than 1 dB. This slight increase was due to irregular evoked signal components not related to white noise, i.e., being related to variation in individual repetitions (amplitude variations, activation jitter and drift).

**Figure 3:**
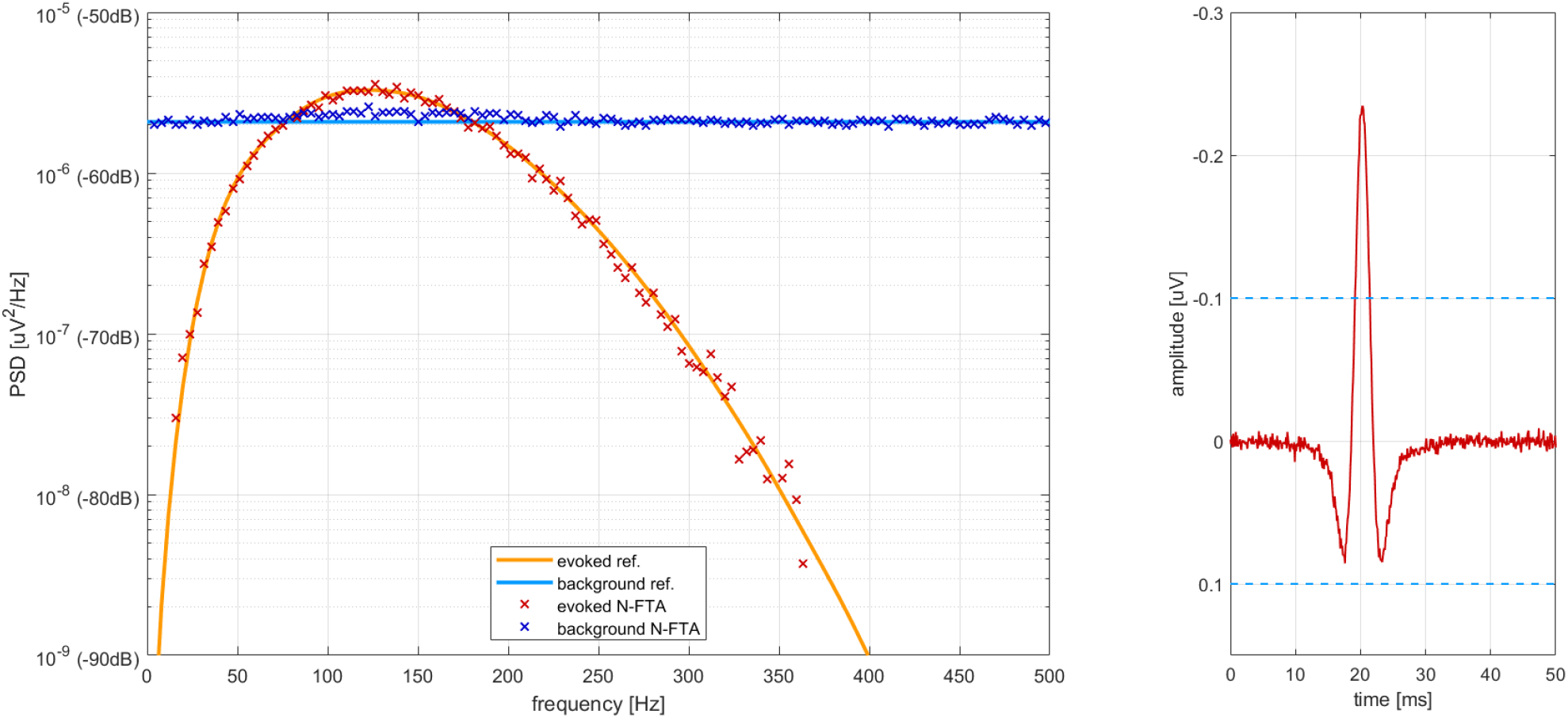
*Left:* Comparison of the *N*-FTA with the reference spectra for the synthetic test data. *Right:* The compound action potential obtained by averaging 1000 responses of the synthetic test data. The dashed lines indicate ± standard deviation of the synthetic white noise in the raw data.

### 3.2. Cortical Median Nerve SEPs

For both subjects the cortical response of highest amplitude was observed for the C3’-Fz channel. Therefore, all presented data was obtained from this derivation.

#### 3.2.1. Low and Medium Frequency Band

Figure 4 shows an example for an amplitude spectrum obtained by a *N*-FT in subject A. In the depicted portion of the MF band (10 Hz to 120 Hz), amplitudes at the evoked harmonics mostly exceeded background activity. In the low frequency band, background amplitude increased significantly, thereby partially covering superimposed evoked harmonics. A high amplitude 50 Hz power line interference was observed, but distinct from the evoked harmonics.

**Figure 4:**
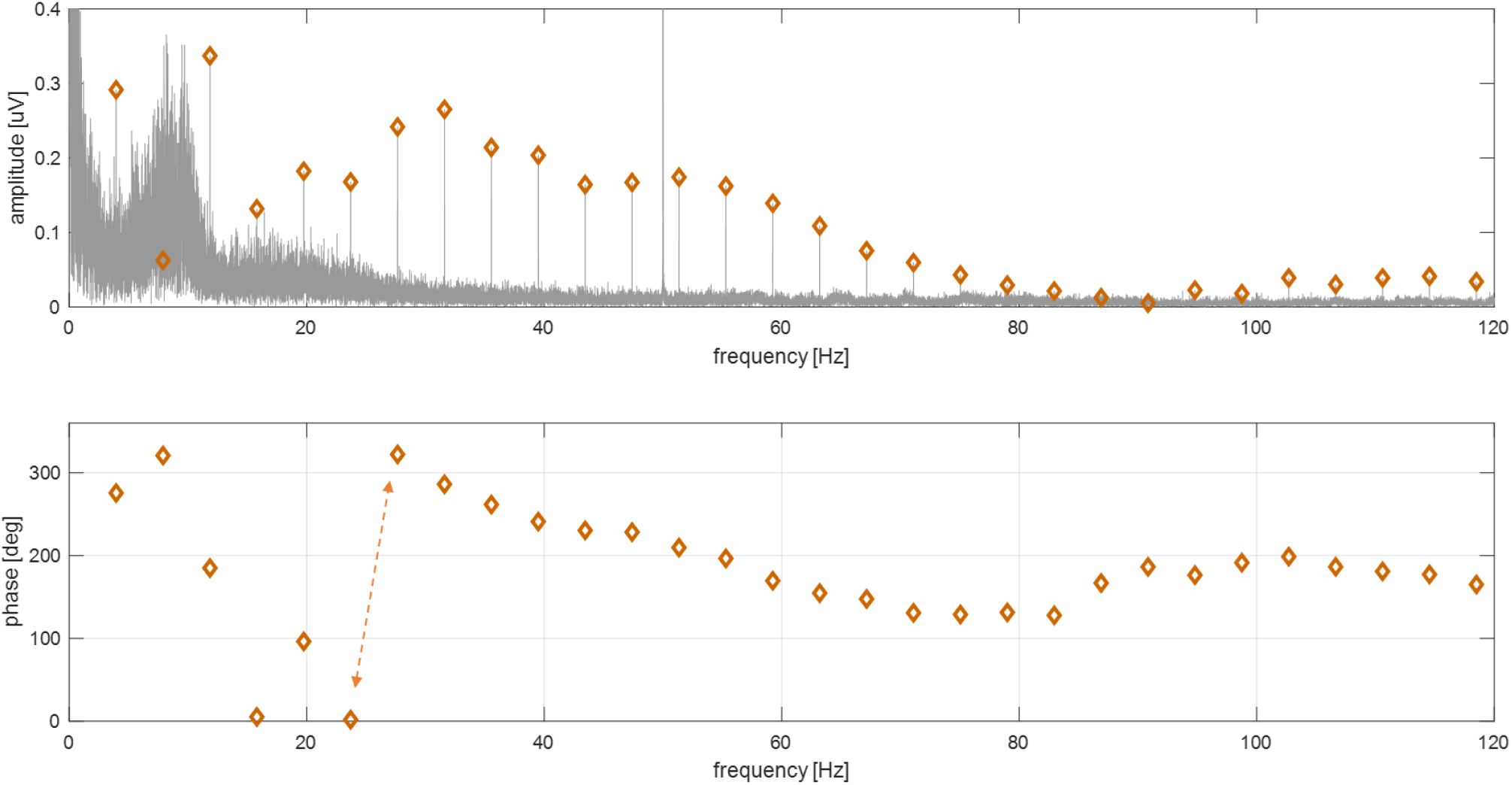
*Top: N*-FT amplitude spectrum for median nerve stimulation at 3.95 Hz (subject A, closed eyes). The interval contained 1009 repetitions (approximately 250 s). The amplitude spectrum is plotted in a gray shade. At the evoked harmonics, diamond markers are displayed. Between 10 Hz to 80 Hz the evoked harmonics clearly exceeded the background activity. Near 10 Hz background activity displayed a peak in the *α* band. At 50 Hz power-line interference was observable. Highest background components occured at near DC frequencies. *Bottom: N*-FT phase at the evoked harmonics for the same trial (time zero 20 ms after the stimulus). Between 50 Hz to 120 Hz phase is near 180° which reflects the N20 peak in the SEP (see also Appendix A.1). The dashed double arrow highlights a 360° phase step (0° to 360° phase interval).

Figure 5 depicts the separation of the data in Figure 4 into evoked and background activity by *N*-FTA. The PSD representation reflects the average power levels for a single response. Thus, the evoked signal displayed a smaller amplitude compared to the background activity. The evoked *N*-FTA data was not continuous since at certain frequencies the evoked activity was strongly interfered by background activity and the acceptance criteria were not met at these frequencies. For example, near the *α*-band and near 90 Hz gaps were observed, which coincided with evoked harmonics below the background level in Figure 4. At frequencies of a few tens of Hz the distance between evoked and background activity was smallest. Within the 100 Hz to 250 Hz interval evoked activity was detected continuously and approximately 20 dB below the background level. Above 250 Hz the evoked activity was often more than 20 dB below the background level and gaps occurred in the evoked *N*-FTA data.

**Figure 5:**
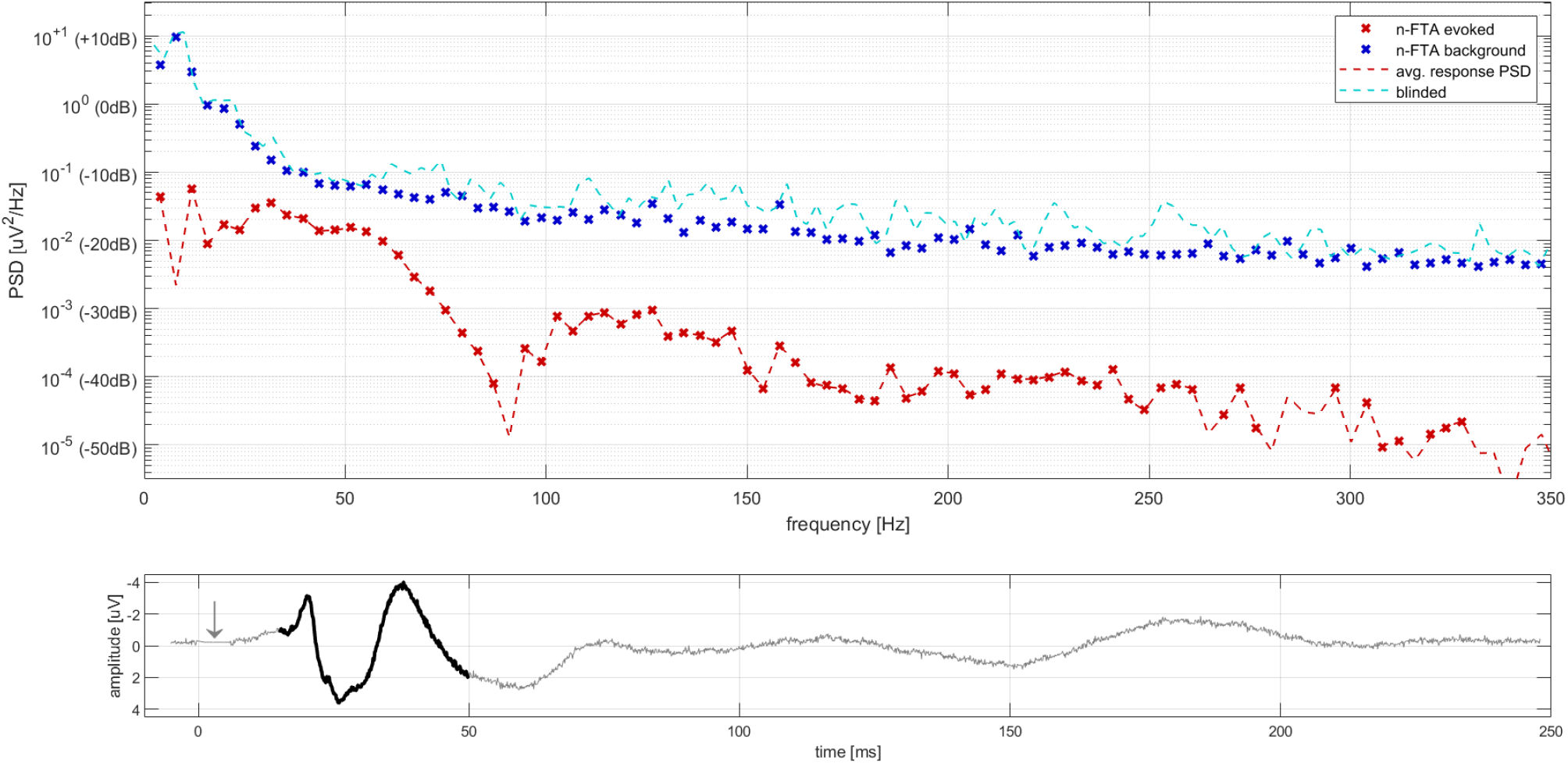
*Top:* Power spectral density plots of evoked and background activity (same trial as in Figure 4). The bold traces display evoked activity (red markers) and background activity (blue markers) obtained from *N*-FTA. Evoked data which failed meeting the acceptance criteria is not shown (see text). For verification background activity was also estimated from the single blinded recording (dashed cyan trace). Evoked activity was verified by the PSD of the averaged response (dashed red line, see text). *Bottom:* The signal as obtained by response averaging of the unfiltered repetitions. Stimulation artifacts and linear drift were removed before averaging (see Figure 2). The arrow marks the signal segment were the stimulation artifact was removed. The diagnostically relevant window was highlighted in black.

In both subjects, the highest background activity was always observed below 6 Hz and at such low frequencies evoked spectral levels were always more than 15 dB below the background components. In the MF band, up to approximately 40 Hz the background level decreased quickly with increasing frequency while further above, a smaller decay was observed. When recording during eyes open the highest background frequency was always at the evoked frequency (i.e., at the lowest frequency captured by *N*-FTA) and approximately 5 dB to 20 dB above the background level obtained during eyes closed at the evoked rate. We observed local peaks in background activity in the *α*- and *β*-bands. The variation in background PSD when recording with eyes open versus eyes closed was in the order of a few dB.

For verifying the background components obtained by *N*-FTA, the PSD obtained from the single blinded recording is also shown in the same plot. Only some minutes passed between the two measurements. Both traces were qualitatively comparable and matched below 50 Hz. At higher frequencies traces run approximately in parallel but for the blind recording PSD showed larger variations. For stimulation at 2.46 Hz deviation between the background levels obtained by *N*-FTA and the blinded recording was 2.8 ± 1.5 dB (mean ± standard deviation) and 1.8 ± 1.0 dB for subjects A and B respectively. The two deviations for the blinded recording with eyes open versus eyes closed were −1.1 ± 2.1 dB and 1.5 ± 1.7 dB for the two subjects, respectively.

For verifying the evoked components obtained by *N*-FTA, the PSD of the averaged but unfiltered response is also shown in the same plot. For evoked components which were accepted by *N*-FTA, both spectral estimators closely matched. The PSD of the averaged response “bridged the gaps” in the *N*-FTA trace. However, in these gaps signal levels were approximately a factor of 1000 (or 30 dB) below the background activity. This factor was caused by the number of stimuli in the data, defining the reduction of background activity by averaging. Thus, the traces in the gaps did not represent an accurate estimation of evoked levels, but reflected essentially the reduction of the background level obtained from averaging.

The separation of evoked and background activity by the *N*-FTA allowed for assessment of EBR – a frequency dependent signal-to-noise ratio. Figure 6 depicts EBR over frequency for 2.46 Hz stimulation and the corresponding SEPs obtained from the medium band setting. The traces displayed broad and flat maxima with some intersubject variation, which was linked to signal morphology. In both subjects, activation of the somatosensory cortex was reflected by an N20 peak of approximately 2 µV amplitude.

**Figure 6:**
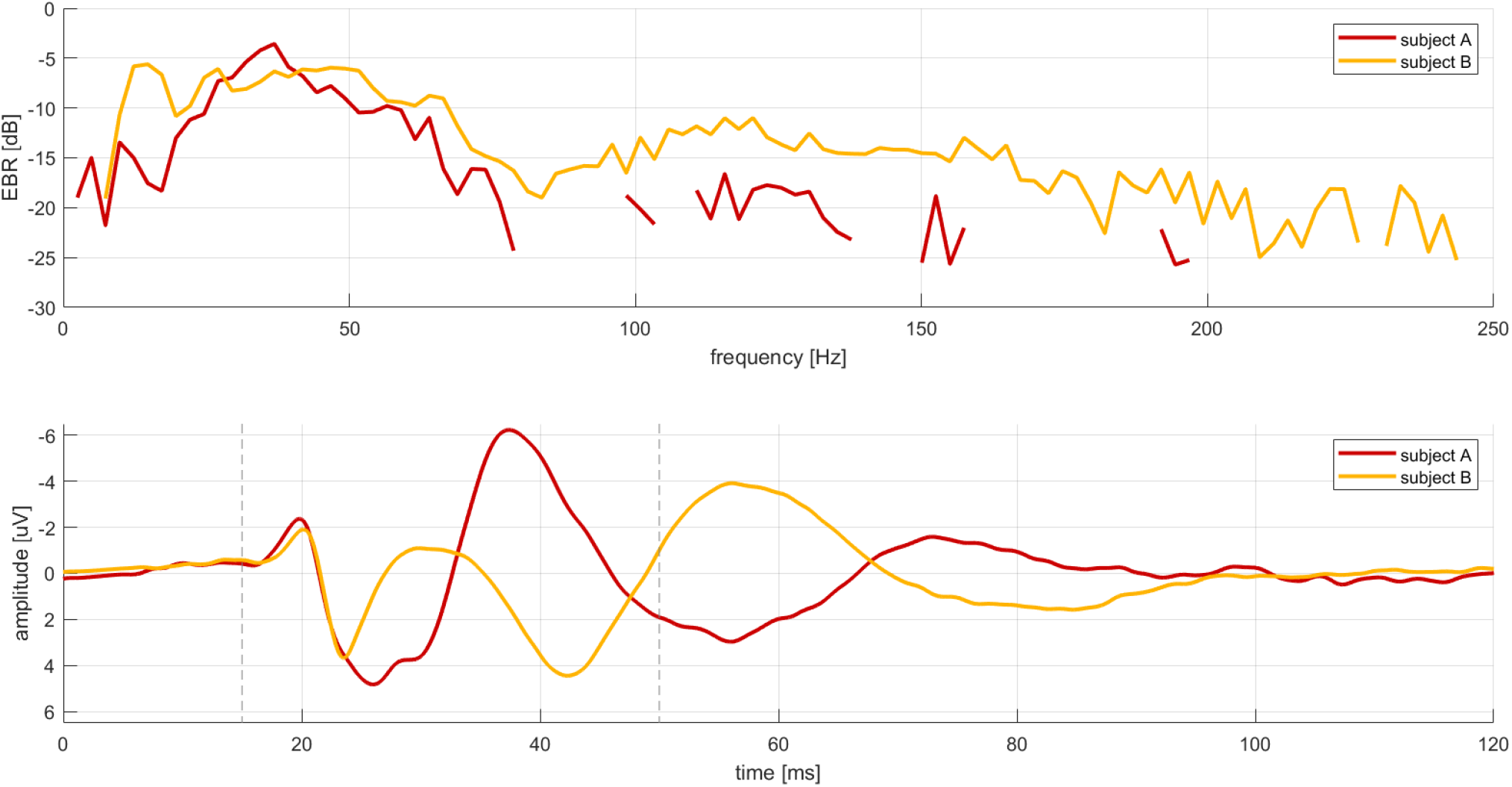
*Top:* Evoked-to-background ratio EBR as a function of frequency for median nerve stimulation at 2.46 hertz in both subjects. *Bottom:* The median nerve SEPs as obtained from the same data by applying the zero-phase filter setting. The diagnostically relevant window was marked by vertical dashed lines. Negativity was plotted upwards.

A broad spectrum was observed for subject B. Here, the interpeak interval between N20 and P25 was 3.4 ms and represented the fastest deflection in the signal. This transition from N20 to P25 may be interpreted as a half cycle of an oscillation of frequency 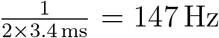. The following half cycles continuously increased in duration. The last half cycles lasted 29 ms (extrema at 56 ms and 85 ms) corresponding to an oscillation frequency of 17 Hz. Thus, SEP morphology of subject B was clearly linked to the broad spectrum, as obtained from *N*-FTA. Also in subject A the fasted transition between two extrema was the N20-P25 segment. However, the P25 peak was broader which was reflected by a lower EBR above 100 Hz. In both subjects fast oscillations were observed within the diagnostically relevant window while in the later phase slower oscillations occurred. Maximum overall peak-to-peak amplitude was 11.0 µV for subject A and 8.3 µV for subject B, respectively.

In the MF band the EBR segment of highest frequency obtained at *f*_*E*_ = 2.46 Hz was found for subject B and contained five evoked harmonics within the interval 231 Hz to 244 Hz. At *f*_*E*_ = 3.95 Hz the highest MF band evoked component was found at 328 Hz. We did not detect any evoked components between 330 Hz and 500 Hz in any trial.

#### 3.2.2. Short Latency SEPs

Figures 7 and 8 show the SEPs obtained from subject A and B for stimulation of the right median nerve, respectively. In the traces which were obtained from the standard setting, high frequency components (a superposition of noise and – near N20 – HFOs) are visible. Furthermore, they contained a pronounced stimulation artifact beginning at time zero. Ahead of the stimulus and in the segment between the stimulus and the onset of the N20 deflection the traces displayed a remarkable near DC offset. The signals obtained from the MF band setting appear smooth. They were near zero ahead of the N20 onset. The stimulation artifact is not visible due to signal pre-processing. Despite the significantly smaller bandwidth of the MF band traces, both settings displayed comparable signal morphology and no relevant systematic time shifts were observable.

**Figure 7:**
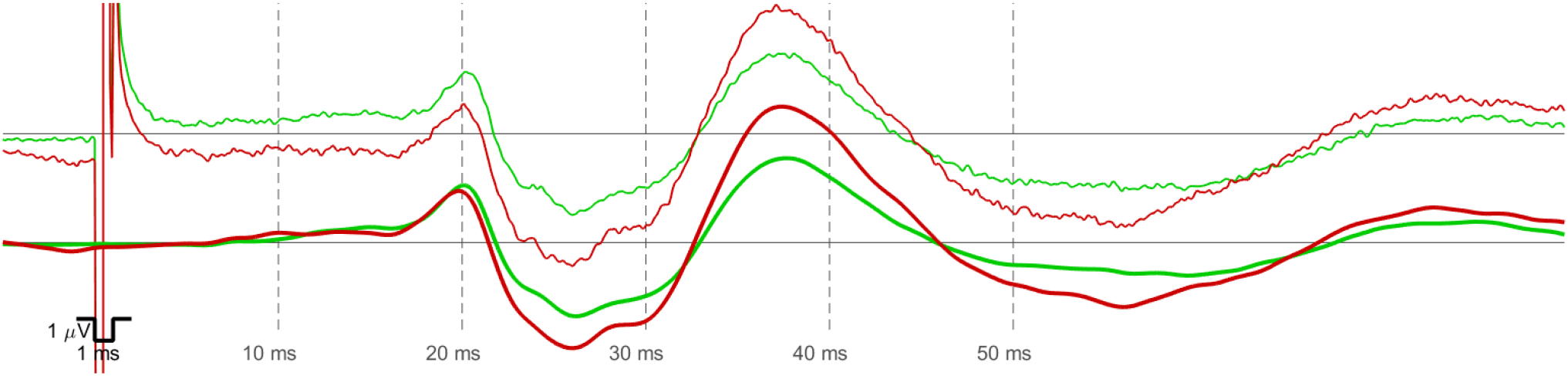
SEPs obtained for stimulation of the right median nerve in subject A. The upper traces show the result obtained from the standard filter setting. Note the offset in isoelectric segments due to the low high-pass corner frequency in the standard setting. The lower traces were obtained from the MF-band setting. The red traces were obtained when stimulating at 2.46 Hz (eyes open). The lower red trace is also depicted in Figure 6. The green traces were obtained at 3.95 Hz (eyes closed). Negativity was plotted upwards as indicated by the calibration pulse.

**Figure 8:**
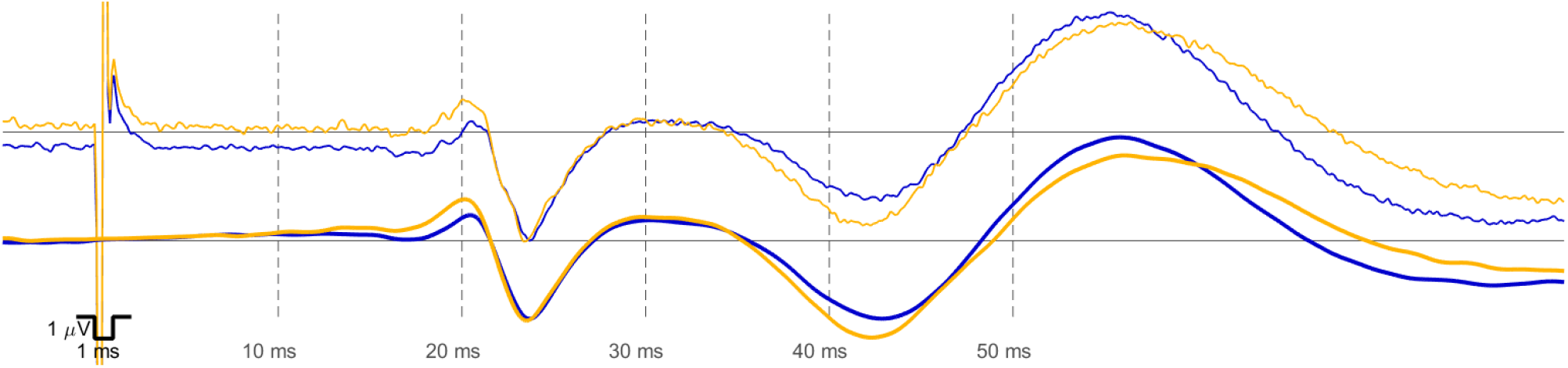
SEPs obtained for stimulation of the right median nerve in subject B. The upper traces show the result obtained from the standard filter setting. The lower traces were obtained from the MF-band setting. The orange traces were obtained when stimulating at 2.46 Hz (eyes open). The lower orange trace is also depicted in Figure 6. The blue traces were obtained at 3.95 Hz (eyes closed). Negativity was plotted upwards as indicated by the calibration pulse.

Table 1 lists data for a quantitative comparison of both settings. The similarity in signal morphology between the settings was reflected by large correlation coefficients (0.90 to 0.99). The difference in N20 latency between both settings was within the range of −0.1 ms to 0.3 ms). For P25 the difference was within the range of −0.1 ms to 0.2 ms). The estimated *SNR*_*r*_ was about twice as high for the MF band setting as compared to the standard setting.

**Table 1:** Correlation, latency and peak-to-peak amplitude for median nerve stimulation.

| subject | eyes | $f_E$ | $N$ | $r$ | setting | N20 | N25 | $V_{pp}$ | $SNR_r$ |
| --- | --- | --- | --- | --- | --- | --- | --- | --- | --- |
| | | Hz | | | | ms | ms | $\mu V$ | |
| A | closed | 3.95 | 1009 | 0.98 | standard | 20.2 | 25.9 | 6.58 | – |
|  |  |  |  |  | MF band | 20.1 | 26.1 | 6.01 | – |
|  | open | 2.46 | 607 | 0.99 | standard | 20.1 | 26.0 | 7.46 | 9.5 |
|  |  |  |  |  | MF band | 19.8 | 26.0 | 7.22 | 17.5 |
| B | closed | 3.95 | 1006 | 0.90 | standard | 20.5 | 23.6 | 5.50 | – |
|  |  |  |  |  | MF band | 20.4 | 23.6 | 4.77 | – |
|  | open | 2.46 | 606 | 0.95 | standard | 20.0 | 23.4 | 6.55 | 7.1 |
|  |  |  |  |  | MF band | 20.1 | 23.5 | 5.59 | 14.7 |

#### 3.2.3. High Frequency Oscillations

For subjects A and B we obtained HFOs near 20 ms following median nerve stimulation at both stimulation rates with a signal level clearly exceeding the background level. The HFOs obtained in subject B for stimulation at 3.95 Hz are depicted in Figure 9. Here, the amplitude of the near N20 oscillation was more than three times the background level. In the 5 ms segment after the stimulus, no oscillations were observable, due to the cancellation of the stimulation artifact. For this trial, *N*-FTA detected evoked components between 517 Hz and 577 Hz (Figure 9 B). They were at a comparable power level as estimated in Appendix C.2.

**Figure 9:**
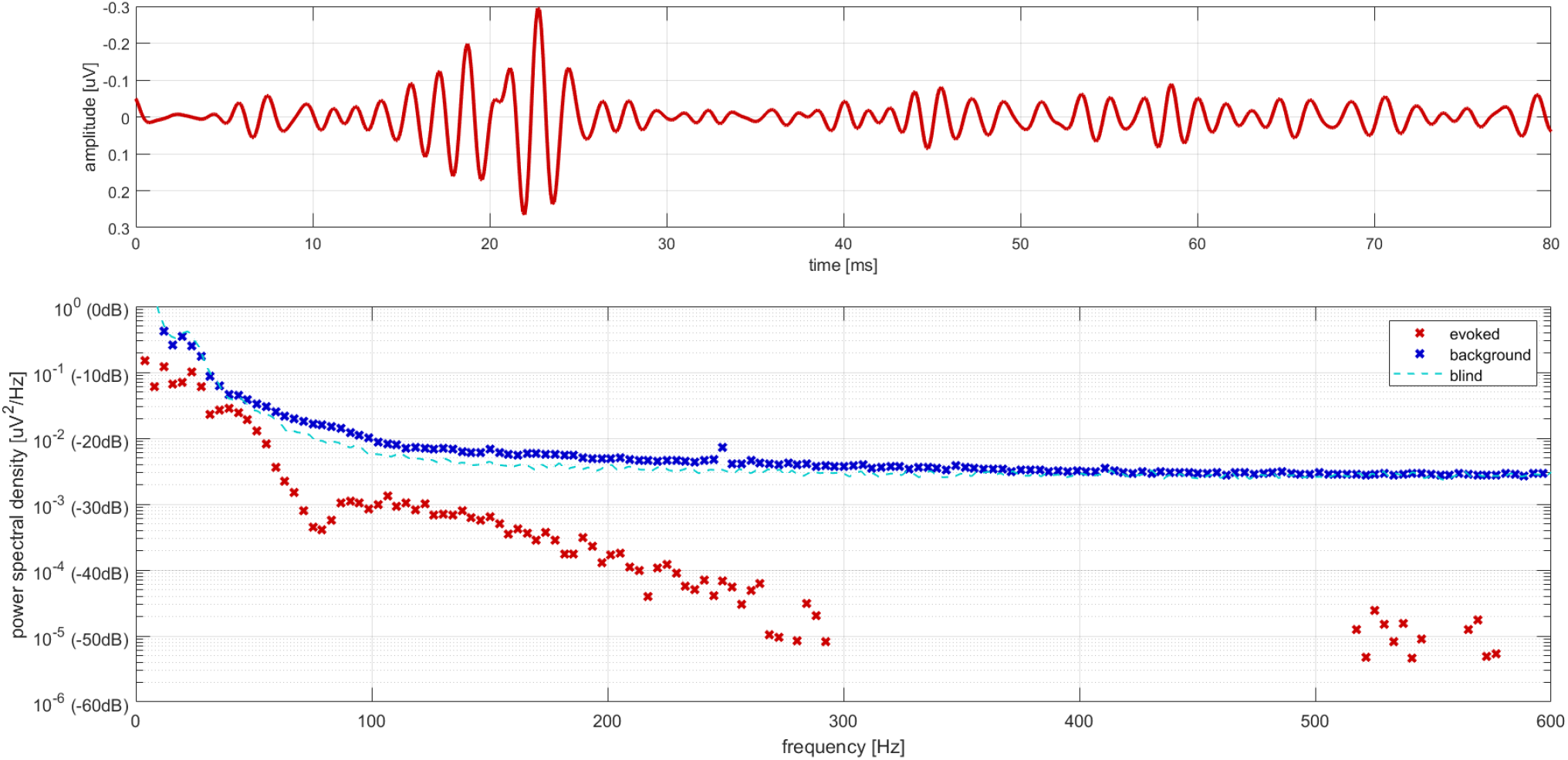
*Top:* HF response to right median nerve stimulation in subject B (lead C3’-Fz, *f*_*E*_ = 3.95 Hz, *N* = 1006). *Bottom:* The evoked and background PSD-components as obtained from *N*-FT Analysis are plotted together with the spectral estimate as obtained from the single blinded recording. High-frequency components were detected in the evoked response.

The evoked spectral components shown in Figure 9 display the longest portion of evoked activity which was detected in the HF-band (450 Hz to 750 Hz). For stimulation at 2.46 Hz in subject B, *N*-FTA detected three distinct segments of evoked activity (each of them by the APSC 2, highest frequency 738 Hz). For subject A at both stimulation frequencies a single HF-segment was detected (again, both by APSC 2).

## 4. Discussion

We developed a novel *N*-FTA approach, which allows for simultaneous spectral investigation of evoked and background activity in individual cortical SEP trials. This enables identification of evoked spectral components being extracted from the raw data with high reliability (i.e. components of a relatively high evoked-to-background ratio, EBR). We aimed to show that a significant portion of diagnostically relevant information (morphology, latency and amplitude of SEPs) is contained within a frequency band which is an order of magnitude narrower as compared to the broad bands which are recommended in clinical guidelines [3, 16]. We presented a signal processing approach which accurately extracts diagnostically relevant parameters only from the MF band (10 Hz to 300 Hz) and compared results with a standard broad band setting.

### 4.1. Synthetic Data

We investigated the accuracy and robustness of *N*-FTA by using synthetic test data. *N*-FTA provides the evoked and background power spectral density at a frequency resolution which is defined by the stimulation rate (evoked frequency *f*_*E*_). At each spectral sample, background activity is the average power level of irregular activity within an interval of width *f*_*E*_. The power of evoked activity gets normalized to that of a single response (thus, corresponding also to a frequency resolution of *f*_*E*_). Therefore, evoked and background levels can be directly compared to each other.

For the synthetic data being corrupted with white noise, evoked activity was detected at sufficiently high power levels. At power levels which were approximately a factor of *N* or more below the noise level evoked activity was not captured. This can be explained as follows: with increasing *N* the spectral resolution in the DFT (the first step in our analysis) increases, which causes a linear increase of PSD at the evoked harmonics with increasing *N*, while the background PSD remains constant for a given noise distribution. As a consequence, *N* determines the threshold for separating evoked and background activity.

In addition to noise, variations in individual cycles were simulated in the test data. These variations caused a slight increase (less than 1 dB) in the *N*-FTA background activity near the peak frequency in the evoked spectrum. This can be explained by variations between cycles that create spectral components that are not related to the evoked harmonics. Thus, inter-cycle variations can be considered as an irregularity properly reflected in background components. Since the effect is caused by an irregularity of the evoked signals it is most pronounced near the evoked peak frequency. Despite the relatively high levels of irregularity chosen for the simulated data (activation jitter and drift were selected at the upper boarder of what was expected for an experimental setting as outlined in Appendix A.4 and amplitude variation was ±50%) inter cycle variation had no significant impact on accuracy, which was reflected by mean deviations below 0.1 dB for evoked and background activity. Consistently with the findings from [11] the CAP spectrum was restricted two a relatively narrow band of a few 100 Hz in width.

### 4.2. Cortical Median Nerve SEPs

#### 4.2.1. Cortical Background Activity

The spectral pattern of background activity as obtained from *N*-FTA and analysis of blinded recordings (without actual stimulation) were qualitatively comparable with other studies on cortical frequency bands [18, 19]. The most pronounced spectral feature was a decay in background activity with increasing frequency which corresponds to the 1/f-like activity described by Voytek et al. Superimposed on this 1/f-like decay characteristic EEG bands like the *α*- or *β*-activity were demarked by local maxima. The variation in background between individual trials and recording with eyes open or closed was in the order of some dB only. At very low frequencies the background PSD increased by more than 10 dB when recording with eyes open. Most likely this was due to eye-blinks, containing frequency components from DC up to approximately 5 Hz [20].

#### 4.2.2. Frequency Bands of Short Latency SEPs

##### Low Frequency Band

Within the LF-band (below 10 Hz) the EBR was relatively low (typically below −15 dB). Thus, evoked LF activity was approximately a factor of ten in amplitude below LF background activity. This was mainly caused by the high background PSD levels which occur in this band (1/f-activity, *θ*−activity in awake volunteers and eye-blinks). The evoked components detected in the LF-band were still of relatively large amplitude. However, as can be seen from the SEPs these slow oscillations occur relatively late after the stimulus and are out of the diagnostically relevant window (15 ms to 50 ms). Thus, the LF-band appears to be of minor relevance for the diagnosis of short latency EPs. The duration of the diagnostic window corresponds to an oscillation of approximately 14 Hz. Obviously, slower signals do not contribute significantly to the morphology of the short latency cortical SEP. Frequencies within the LF band might potentially be of relevance for the diagnostic use of slow responses at stimulation rates ≤ 0.5 Hz [21].

##### Medium Frequency Band

Within the 10 Hz to 300 Hz MF-band we identified broad segments of relatively high evoked activity. The largest EBR was near −5 dB (i.e., evoked activity being approximately a factor of two in amplitude smaller compared to background activity) and was always found at some tens of Hz. In one subject we observed a relatively high EBR between 100 Hz to 200 Hz and we were able to link this spectral portion to a short N20-P25 interpeak interval (3.4 ms). Also for the other subject we were able to identify a long segment of evoked activity above 100 Hz (Figure 5). These broad MF segments of evoked activity significantly contribute to signal morphology in the diagnostically relevant window. Thus, a significant portion of diagnostically relevant signal features such as morphology, latency and amplitude are contained in the MF band. We did not detect any evoked activity between 350 Hz to 500 Hz, consistent with only rather small evoked signal levels at such frequencies.

##### High Frequency Oscillations

We investigated the HF-band between 450 Hz to 750 Hz. At these high frequencies the detection of evoked activity by *N*-FTA approaches its limits due to the following reasons: firstly, evoked HF-components are small (about −30 dB to −20 dB below background activity) and, secondly, small variations in individual responses may reduce the sensitivity at several 100 Hz (see Appendix A.4). In spite of this, *N*-FTA detected at least one segment of HF-components in each trial and multiple segments in two trials. Considering the low threshold which was chosen for accepting false positive evoked components, it is highly probalble that the algorithm captured true HFO-components near the boarder of the obtainable resolution. The assessment of background activity was not limited in the HF-band.

To confirm the presences of HFOs in the signals we averaged bandpass filtered HF-data identically as described in [6]. For all trials we obtained HF-oscillations near 20 ms with peak-to-peak amplitudes of some tenths of µV. This was comparable to other studies [6, 5]. However, these low HF-amplitudes are about an order of magnitude smaller as the short latency amplitudes of standard SEPs. Thus, the HFOs may have only a minor influence on the N20 peak latency.

#### 4.2.3. Short Latency SEPs

As outlined in the introduction, clinical SEP settings use broad filter bands for historical reasons. As a consequence, routinely recorded signals appear almost unfiltered, which affects routine usability and robustness negatively.

##### Signal Features

We showed in two subjects that key features of cortical short latency median nerve SEPs can be accurately obtained from a restricted MF-band. Firstly, we reduced the low-pass corner frequency to 240 Hz, which was about a factor of ten smaller than the clinical standard (1 kHz to 3 kHz [3, 16]). Secondly, we increased the high-pass corner frequency to 18 Hz, which was again about an order of magnitude above the clinical standard (1 Hz to 5 Hz). We compared the medium band setting to a standard setting. For all four trials, the MF-band setting was able to properly reflect signal morphology and to accurately capture latency, despite the drastic reduction in signal bandwidth. The agreement in signal morphology was reflected by a high correlation of 0.90 to 0.99 within the diagnostically relevant window and deviations in latency were in the order of a few tenths of ms which is an accepted limit for clinical reproducibility of latency measurements [3]. Interestingly, the deviations in latency between settings was much smaller than the action potential duration of a single neuron. Furthermore, amplitudes were comparable for both settings but slightly larger for the standard setting. This appears to be due to HF-components (evoked activity and noise) contributing to a slight increase in peak-to-peak amplitude. However, at least the observations from two subjects suggest, that the removal of HF-components in the MF-setting still allows for extraction of diagnostically relevant features with sufficient accuracy.

##### Filter Concept

The accurate assessment of latency from a narrow MF-band became possible due to the use of two techniques: firstly, the use of zero-phase filters and, secondly, the removal of stimulation artifacts. The basic theory of zero-phase filters was established decades ago. With increasing use of digital signal processing the continuously spread into many applications. However, their ability to accurately maintain latency at reduced bandwidth was perhaps overlooked. Even some recent studies use wide band filtering and analog or digital filters generating phase shifts [17]. Recent advances in biopotential amplification and AD-conversion technology allow for development of novel filtering concepts [2]. State of the art 24-bit A/D conversion devices allow for avoiding all analog filters and for tailoring digital signal processing for suppression of different types of artifacts with different filter types. Stimuli can be removed by using time domain subtraction and zero-phase filters can be accurately tailored to the frequency band of interest.

##### Signal-To-Noise Ratio

For the MF-band setting, twice as large SNR estimates were obtained, reflecting a high MF-band EBR, while the broad band of the standard setting contained large portions of poor EBR. For the standard setting, in particular the high amplitude LF-background activity (see above), is only partially suppressed by averaging. This remaining LF interference appears as offset in the traces (see Figures 7 and 8) but does not reflect any short latency potential. The MF band setting filters LF interference out (in addition to averaging), thereby reducing the remaining noise floor.

### 4.3. Future Developments

#### N-FTA

One scope of this study was to establish the methodological background for simultaneous assessment of evoked and background activity in individual trials. Future work should broaden this application. Significantly larger cohorts of individuals will allow for statistical assessment of EBR distributions and eventually investigation of systematic changes in spectral profiles caused by various diseases. Other stimulation sites (e.g. tibial or sural nerve) or different signal types (e.g., travelling near-field potentials in peripheral nerves or spinal tracts, stationary far-field potentials recorded from spinal or scalp positions) are potential targets for scientific investigation. We chose median nerve short latency cortical SEPs for methodological development, since it is a signal of high diagnostic importance of which a large body of literature (including investigations of the HF-band) exists. In addition to short latency SEPs also late evoked responses (at stimulation rates ≤ 0.5 Hz) or visual, auditory or tactile evoked potentials can be studied in future trials.

#### SEP Recordings

Another scope of this study was to show the possibility to extract diagnostically relevant features of short latency cortical SEPs from an MF-band which is an order of magnitude more narrow as compared to standard diagnostic settings. In a first step, analysis was performed offline. Routine application, however, requires real time signal processing. Thus, methods are required for online removal of stimulation artifact and zero-phase filters need to be developed which can work with temporal delays which are acceptably short. Such systems being able to extract the diagnostically relevant signal features from a relatively narrow MF-band may ease and speed up application in clinical settings, since they are potentially less prone to interference. The reliable estimation of signal quality from individual recordings might further contribute to improved usability of EP recording systems.

## 5. Conclusions and Outlook

We present a novel *N*-FTA approach, which allows for simultaneous spectral investigation of evoked and background activity in individual SEP trials. For short latency median nerve SEPs, evoked-to-background ratio was below approximately −15 dB. This was due to the high amplitude of near DC background activity (1/f-activity, eye blinks, *θ*-band, etc.). Largest EBR of approximately −5 dB was obtained at some tens of Hz. Broad segments of evoked activity were identified within the 10 Hz to 300 Hz MF-band. Within the 450 Hz to 750 Hz evoked HF components were detected approximately −25 dB below the background level at the boarder of the algorithm’s sensitivity. The *N*-FTA approach can be applied in future studies for accurately investigating EBR in different frequency bands of essentially all types of repetitively evoked potentials.

We applied a narrow band filtering approach for obtaining median nerve cortical SEPs from a restricted frequency band and compared results to a standard broad band setting. Diagnostically relevant features (signal morphology, latency and amplitude) can still be extracted with sufficient accuracy from the MF-band. Distinct HF-signal components were clearly detected near 600 Hz but had only minor influence on the recorded SEPs. Algorithms should be developed in the near future, which allow for real time extraction of short latency responses from a relatively narrow MF-band. This might render the technology less prone to inference and might improve robustness and clinical usability.

## Acknowledgments

The authors want to thank Nadia Obex for her assistance in performing the experimental recordings. This work was funded by the Tyrolean Science Fund (contract nr. F.18687).

## Declaration Statements

### Conflicts of Interest

None.

### Ethical approval

The study was approved by the institutional *Research Committee for Scientific Ethical Questions* (RCSEQ 2632/19).

## Online Supplement

This document provides the Appendices for the manuscript submitted to Computational and Mathematical Methods in Medicine.

## Appendices

### A *N*-Interval Fourier Transform Analysis

The scope of this section is to provide detailed information on *N*-Interval Fourier Transform Analysis (*N*-FTA, described in the manuscript). In subsection A.1 we summarize relevant theory from the literature on finite periodic repetitions and illustrate observations by an example. Subsection A.2 describes implementation and highlights important aspects for obtaining sufficient accuracy. Subsection A.3 provides a verification of the accuracy of the implemented code by means of an example. Finally, subsection A.4 investigates the influence on variations in individual responses by reviewing theory and by studying numerical examples.

#### A.1 Finite Periodic Signal – Theory

As a basic model, repetitive evoked recordings may be approximated by a series of *N* identical responses which are corrupted by irregular background activity [4]. Figure A.1 depicts an example for illustration. Here, the evoked signal contained *N* = 4 identical simulated compound action potentials (CAPs, as obtained from [1]). They were repeated at a stimulation interval *T*_*E*_ of 100 ms (i.e. 10 Hz stimulation rate or evoked frequency *f*_*E*_). White noise with a standard deviation of 0.1 µV was used as a model for irregular background activity.

A signal containing *N* identical repetitions is a finite periodic signal. In a strict mathematical terminology it is **a-periodic** as it contains two isoelectric segments of infinite duration. Writing *ℱ*{*ϕ*_1_}for the *time-continuous* Fourier transform of a single response *ϕ*_1_, the FT of a series *ϕ*_*N*_ of *N* identical responses has been obtained from [4]

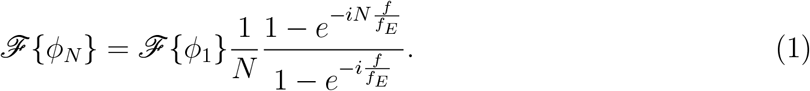

Figure A.1 shows the magnitude of the *time-continuous* Fourier transform (FT) of the exemplary finite periodic signal. FT showed local maxima at multiples *kf*_*E*_ (i.e. at the evoked harmonics) and regularly spaced zero crossings in between the evoked harmonics. This pattern was governed by eq. (1). Experimental data requires the investigation of a finite time interval. For convenience we chose exactly the finite periodic interval. Generally, DFT assumes that the signal was periodically repeated outside of the finite periodic interval. Thus, we obtained a **periodic** signal with a fundamental frequency *f*_*E*_. The discrete frequency samples were exactly at the evoked harmonics or at the zero crossings (Figure A.1). A frequency resolution of 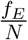 was obtained. We refer to the DFT of a finite periodic interval by the term *N*-FT.

**Figure A.1:**
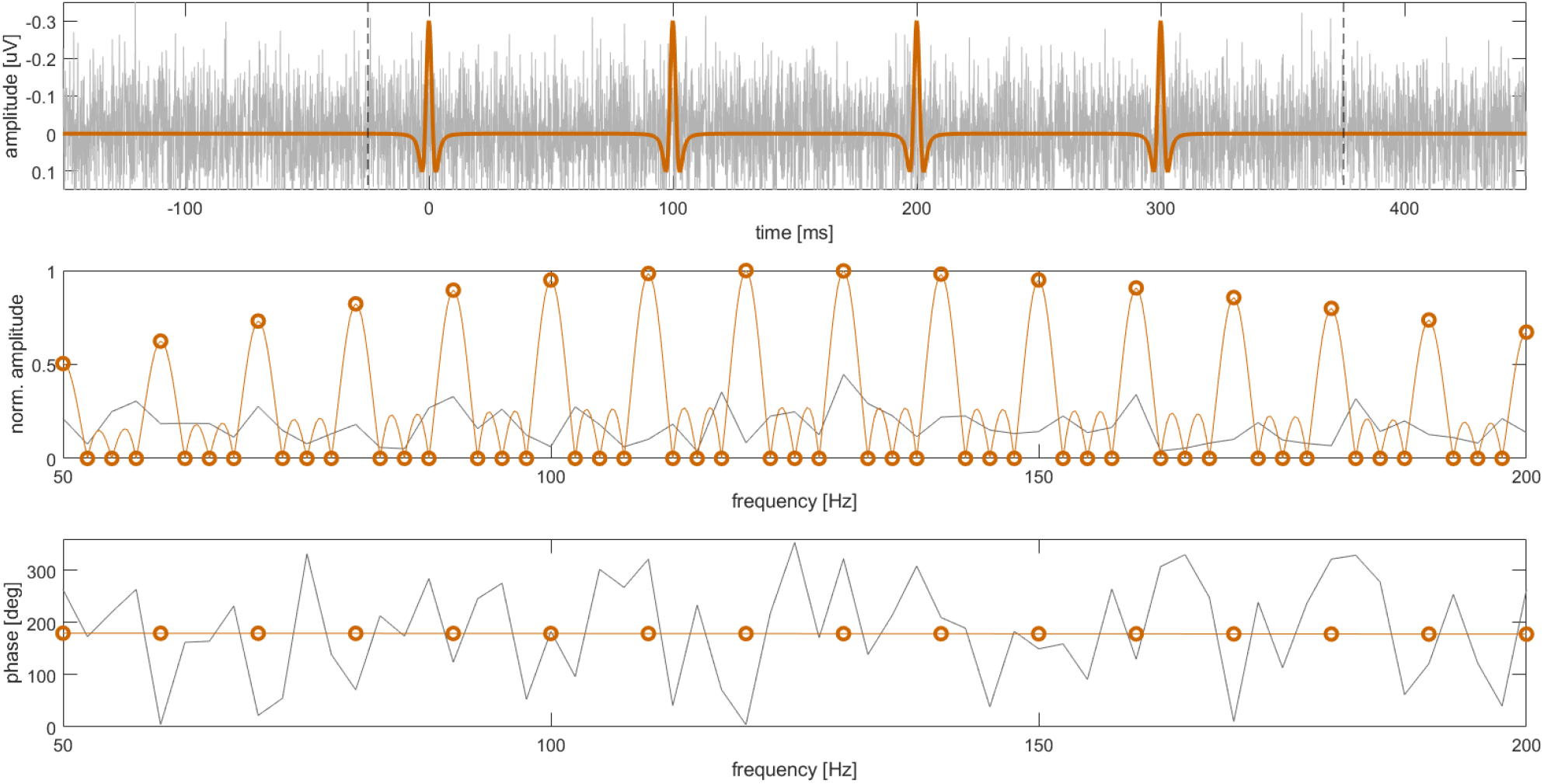
*Top panel:* Finite periodic signal with *N* = 4 repetitions (orange) at *f*_*E*_ = 10 Hz and stochastic white noise (gray). The vertical dashed lines mark the finite periodic interval. According to clinical practice, negativity was plotted upwards. The first peak was centered at time point zero. *Middle panel:* Time-continuous FT of the finite periodic signal (narrow orange line) and its *N*-FT (orange circles) within a frequency interval around the maximum. White noise displays random variations in the spectrum (gray trace). *Bottom panel:* For the finite periodic signal phase was plotted for multiples of *f*_*E*_ (orange markers). For white noise phase was stochastic (gray). The same frequency interval as in the middle panel was used.

As can be observed from Figure A.1, the *N*-FT of the finite periodic evoked signals contained non-zero components only at the evoked harmonics. An irregular background signal contained non-zero elements at all frequencies. We summarize therefore:

- Harmonic components *kf*_*E*_ represent a superposition of the evoked signal and irregular background activity.
- Non-harmonic components represent the spectrum of irregular signals, such as noise or physiological background activity.

Beside amplitude phase was investigated. We observed from the time shift theorem of Fourier transform, that phase depends on the definition of the time zero point. It was reasonable defining time zero such, that it is at (or near) a representative morphological marker in the signal. For the example in Figure A.1, the negative peak defined a representative morphological marker. For the almost even shape of a CAP pulse in the example, the *N*-FT at the evoked harmonics contained essentially cosine terms with a negative sign. Thus, phase was almost constant and near 180°. Since *N*-FT considers a periodic evoked signal, we obtained identical phase for selecting time zero at any peak of the *N* repetitions.

In the experimental part of this study, we chose the N20-peak of a median nerve SEP as the morphological marker. Thus, a phase angle near 180° reflected negative cosine components centered around the N20 peak. We defined time point zero within the first repetition cycle, 20 ms after the first stimulus.

#### A.2 *N*-FTA – Implementation

##### Numerical value of the evoked frequency

In our experimental recordings containing some 100 s of data at a sample rate near 10 kHz, the number of data points *M* in the interval exceeded a million. For accurately capturing the evoked harmonics in the spectrum the evoked frequency must be numerically represented with an accuracy in the ppm (parts per million) range. Note that for state of the art electronic devices, the uncertainty in stimulation frequency or sample rate is in the order of some ppm.

In our software implementation, we chose the following approach for obtaining a sufficiently accurate numerical value for the evoked rate *f*_*E*_. We chose the time frame of data sampling as the reference and set the sample rate *f*_*S*_ = 9.6 kHz (i.e., the nominal sampling rate of the used amplifier). We computed *f*_*E*_ based on this reference. In our experimental data the segment after the last stimulus was not complete. Thus, the data contained *N* + 1 stimuli but *N* usable repetitions. Our code determined *M* as the number of time steps from the first stimulus to the last time step ahead of stimulus *N* + 1. The evoked frequency was then obtained by:

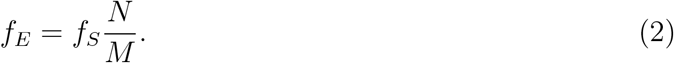

For our experimental data the computed evoked rates were about 17 ppm below the nominal values (2.46 Hz and 3.95 Hz).

###### Pseudocode

*N*-FTA applies three steps. In a first step (see Algorithm 1) the preprocessed signal (i.e., the signal after removal of stimulation artifacts and drift) is loaded and an DFT of the finite periodic interval is performed (*N*-FT). Then, power spectral density *w* and phase information *θ*′are computed (see also manuscript). Here, the function “CheckInterval” ensures that angles are within the interval [0°,360°]. A data structure *St* is constructed containing all harmonic frequencies below the Nyquist frequency. The number of repetitions *N* is stored in the structure *St*.

In the pseudocodes we use the notation ⌊…⌋ for rounding towards lower integer (“floor”), ⌈…⌉ for rounding towards upper integer (“ceil”) and ⌊…⌉ for rounding towards nearest integer (“round”).

In a second step, the background activity is assessed (see Algorithm 2). Within a loop background activity is computed at each evoked harmonic *kf*_*E*_ and the width of the window at each evoked harmonic is slightly below ±*f*_*E*_/2 (“floor” rounding of border indices). The PSD at *kf*_*E*_ was always removed from the analysis, as it contains evoked activity. The detection of power line harmonics is based on the observation, that for a powerline harmonic *nf*_*PL*_, the “check variable” *f*_*check*_ = *kf*_*E*_ − *f*_*PL*_ must be within the interval [−*f*_*E*_/2, *f*_*E*_/2]. A while-loop is used such that *f*_*PL*_ is subtracted exactly *n*-times from *f*_*check*_. If the check variable is within the interval, frequencies near the powerline harmonics (interval ±Δ*f*_*PL*_/2) are removed from the analysis. The average PSD of the data addressed by the indices in the array defined the background level.

###### Algorithm 1 Prepare Data for *N*-FTA

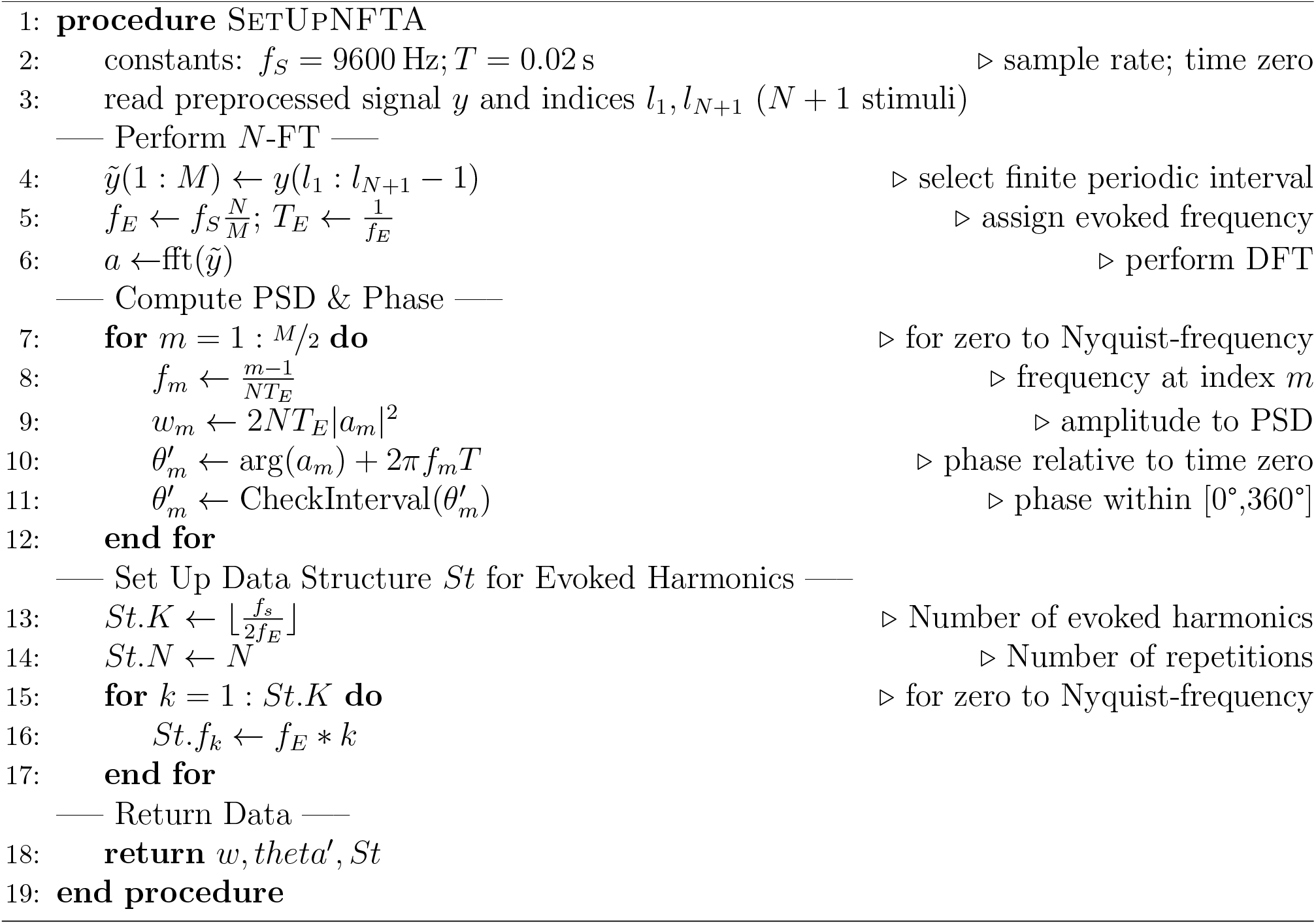

The *array* is also used for computing the percentile when comparing PSD at the evoked frequency with the background level. Note that background activity gets sampled with the frequency resolution *f*_*E*_.

The analysis of evoked frequency components requires a decision about acceptance of each evoked harmonic. The pseudocode in Algorithm 3 first initializes the structure field *St.A* with `0` for all elements. The code then checks for Amplitude Phase Acceptance Criterion (APSC) 1. Upon acceptance of an individual harmonic, the value gets updated to `1`. APSC 2 investigates evoked harmonics together with their left and right neighbors. If all three harmonics in such a group were larger than *p*_*L*_, a phase check (as described in the manuscript) is performed. Here routine CheckPhase selected a proper 360° interval such that no phase steps occur between neighboring harmonics. Upon acceptance *St.A* was set to `1` for all three elements of group. Finally, the PSD *St.e*_*k*_ is computed for all accepted harmonics and normalized to the level of a single repetition. For harmonics, which did not pass the acceptance criteria, the numeric data type `NaN` (not a number) is assigned to *St.e*_*k*_.

As the result of the *N*-FTA the structure *St* contains the evoked frequencies *kf*_*E*_ together with the associated background and evoked PSD levels *b*_*k*_, *e*_*k*_. The evoked frequency *f*_*E*_ defines the frequency resolution of the method. For the presented pseudocode, *St.f*_*k*_ contains frequencies up to the Nyquist frequency. However, in our study we included only frequencies up to 750 Hz into the analysis. This was motivated by a reduced accuracy of the algorithm above 1 kHz, as will become evident from section A.4.

###### Algorithm 2 *N*-FT Background Analysis

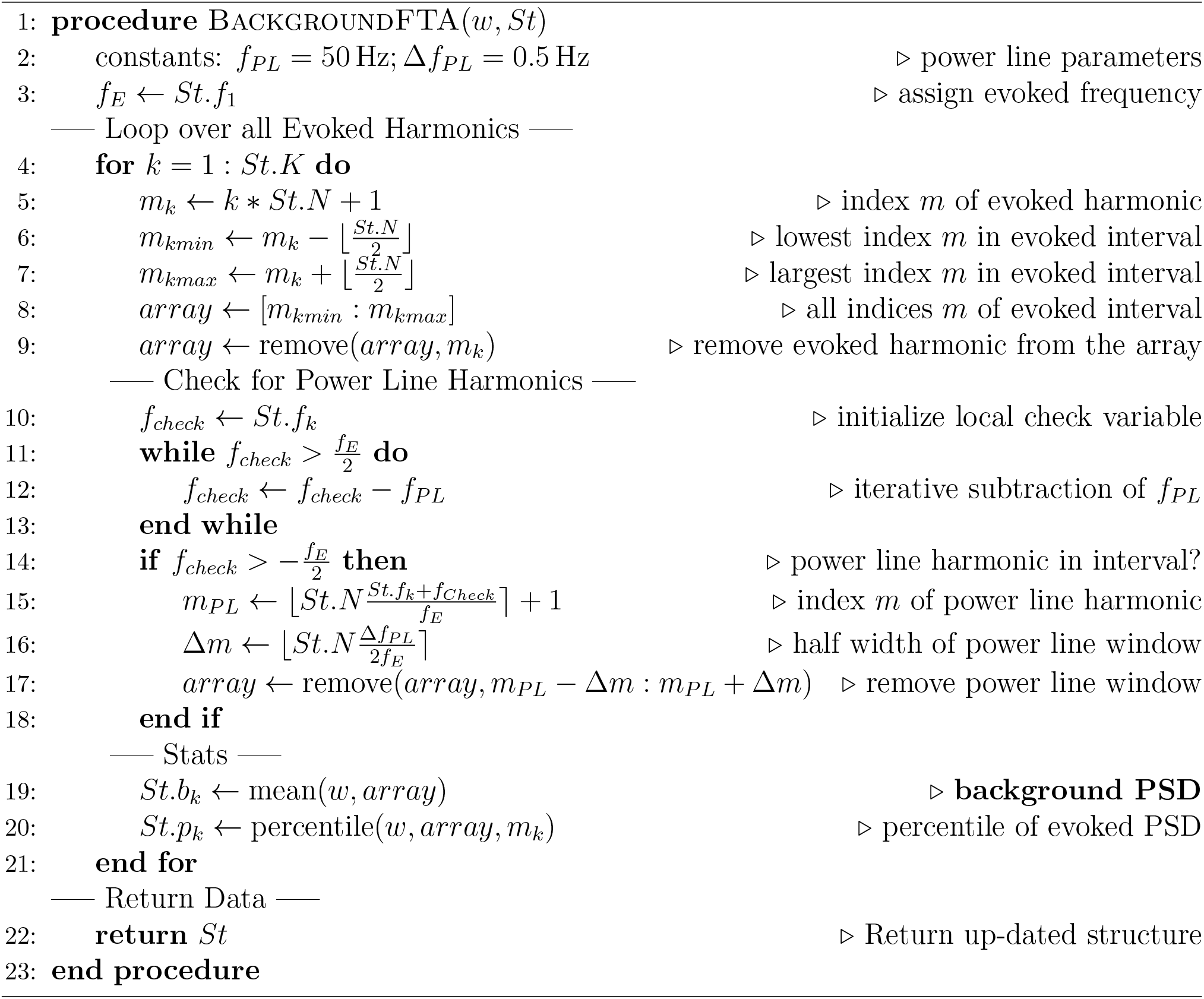

###### Algorithm 3 *N*-FT Evoked Analysis

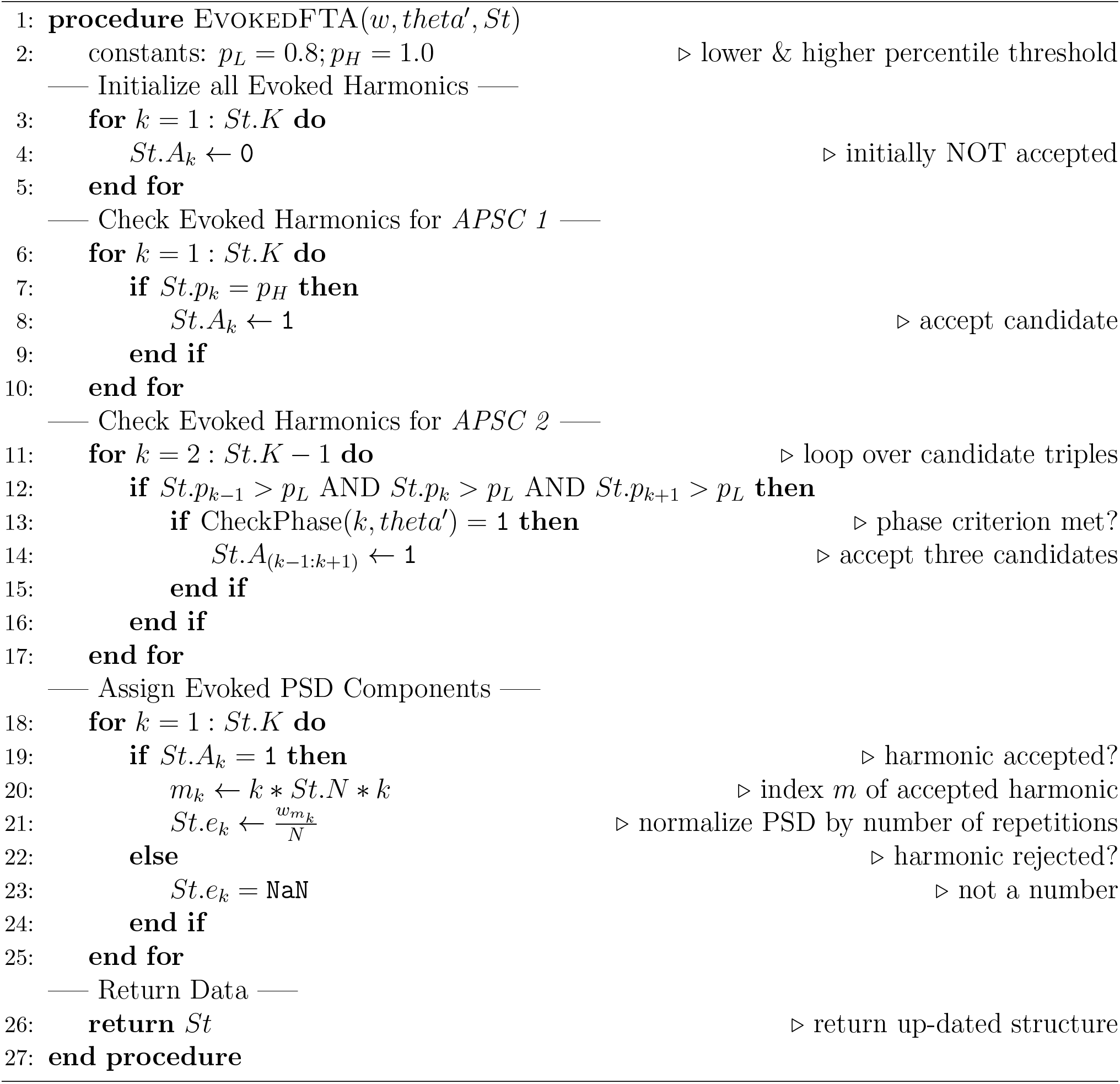

#### A.3 *N*-FTA – Verification

For testing function and accuracy of the implemented code, a numerical example was created. Parameters were chosen such, that they match with the values of the experimental setting, i.e., *N* = 1000, *f*_*E*_ = 3.95 Hz and *f*_*S*_ = 9.6 kHz. As a template for a single response, a simulated CAP was used. The data was taken from a previous study [1] (simulation parameters: conduction velocity *v* = 60 m s^*−*1^, depth of the neural tract *s* = 50 mm and temporal dispersion *τ* = 1.25 ms). This signal and its amplitude spectrum are depicted in Figure A.1. Its peak-to-peak amplitude was 0.32 µV. The original simulation was digitized at a sampling interval of 5 µs. For verification purposes the data was down-sampled to *f*_*S*_ = 9.6 kHz within an interval of length *T*_*E*_. Time zero was chosen such, that the negative peak occurred at 20 ms. For creating a finite periodic evoked CAP this segment was repeated *N* times. The reference PSD for the evoked CAP was obtained by performing a standard DFT for the template of a single response and converting the result to a PSD representation.

For test purposes the signal was corrupted with two levels *σ*_*W*_ of white noise: 0.01 µV and 0.1 µV, respectively. For each noise level *σ*_*W*_ the corresponding background PSD *b*_*W*_ was obtained from

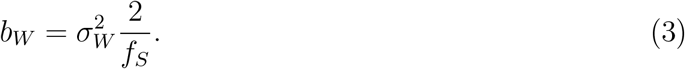

Thus, the two investigated noise levels corresponded to spectral background levels of −56.8 dB and −76.8 dB, respectively (relative to the 1 µV^2^/Hz level).

Figure A.2 depicts the reference PSD trace together with the evoked and background components obtained by *N*-FTA at both noise levels. The two computed background levels match the predicted levels with a residual deviation of 0.00 dB ±0.13 dB (mean ±standard deviation) and 0.09 dB ±0.18 dB at noise levels of −56.8 dB and −76.8 dB, respectively. Note that the standard deviation reflects the remaining random variation for the individual white noise pattern. Consistent with the findings from [1] the evoked spectrum is of a band-type with a peak at 122.5 Hz. For both noise levels the computed evoked levels match the reference well within the frequency band from 50 Hz to 260 Hz. Out of this band small but visible deviations were observed. For the higher noise level the evoked frequency was detected from 12 Hz to 380 Hz. For the lower noise level the interval of detected evoked activity was 4 Hz to 446 Hz. Evoked activity was detected down to approximately 30 dB below the noise level. The errors for the detected evoked components were 1.00 dB ± 1.11 dB and 0.05 dB ± 0.69 dB for the high and low noise levels, respectively.

Thus, we verified that *N*-FTA detects and separates background and evoked activities. The limit for detecting evoked activity is approximately 10 log_10_(*N*) [dB] below the background level. The accuracy of the detected evoked components improves with increasing evoked-to-background ratio.

**Figure A.2:**
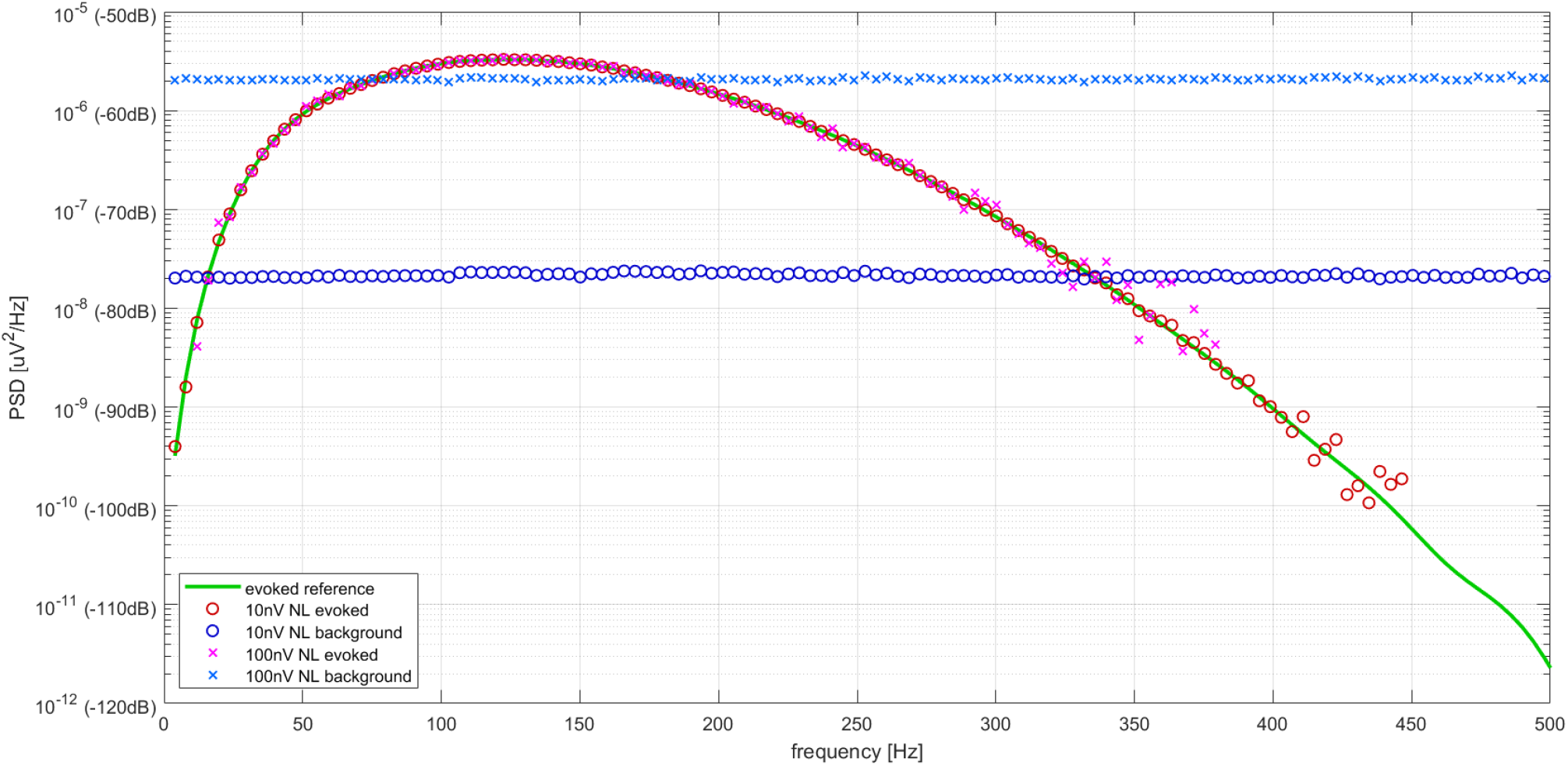
Verification of *N*-FTA at two noise levels (0.01 µV and 0.1 µV) by comparison with a reference evoked activity (see text).

#### A.4 Variations in Repetitions

As it was outlined in section A.1, *N*-FTA assumes that all repetitions were identical. However, in real world experimental settings (small) variations may be observed between individual responses. The scope of this section is to investigate the influence of inter-cycle variations on the accuracy of *N*-FTA. We investigated three different types of variations and quantify their effect.

##### Irregular Activation / Jitter

Random variation of the activation threshold causes irregular activation (or activation jitter). Rompelman and Ros investigated irregular activation in part two of their review [5]. The upper row in Figure A.3 provides a qualitative illustration of their findings. Jitter goes along with a broadening of harmonic peaks at higher frequencies. The work of Rompelman and Ros provides an analytic treatment, which allows for an estimation of the Fourier transform for a given statistic jitter distribution *δ*(*T*). Here, the Fourier transform of the distribution function *ℱ*{*δ*(*T*)}, i.e., the harmonic function of *δ*(*T*), enters the calculation.

As can be seen from Figure A.3 activation jitter has less influence on spectral components of lower frequency. We aim to simplify the analytic expressions listed in [5] such, that a border frequency *f*_*B*_ can be estimated, below which jitter has only a minor influence. For the ideal case of regular activity the distribution *δ*(*T*) becomes a Dirac pulse and its harmonic function is a constant in the frequency domain. As we have shown in the Appendix of [1] the harmonic function of any distribution function is of a “low pass type”, i.e., it is approximately constant at low frequency and approaches zero at higher frequency.

For a normal distribution *δ*_*n*_(*T*) with standard deviation *τ*, the harmonic function is a Gaussian curve with a standard deviation of 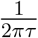. For estimation we assumed that for frequencies below half the standard deviation *τ* the harmonic function was sufficiently close to a constant value and obtained

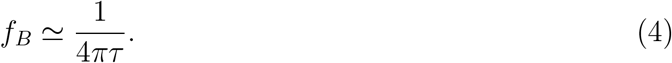

**Figure A.3:**
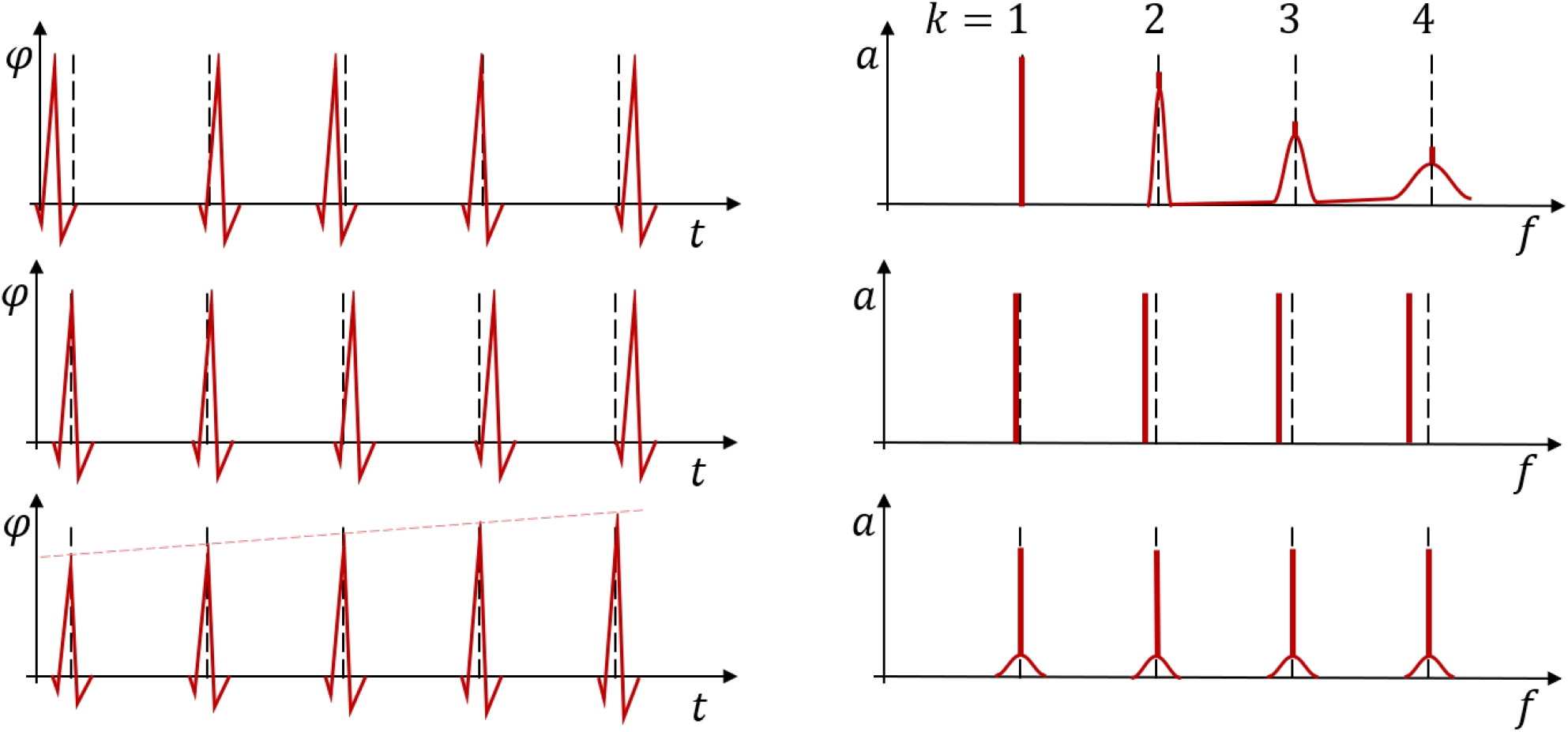
Different types of variations in repeatable signals and their influence on the spectral profile. *Top left:* Irregular activation occurs with a random time shift from a regular action timing (indicated by hashed lines). *Top right:* In the frequency domain irregular activation is reflected by broadening of harmonic peaks with increasing index *k. Middle left:* Activation drift is reflected by a systematic shift in activation timing increasing in magnitude with each repetition. *Middle right:* In the frequency domain activation drift is reflected by a systematic displacement of the harmonic peaks. The magnitude of the displacement increases with index k and its sign it opposite to the time shift. *Bottom left:* A variation in amplitude is reflected by a change of amplitude over time. *Bottom right:* In the frequency spectrum variation of amplitude is reflected by side bands adjacent to each harmonic peak. The peak frequency remains unaffected.

As it was shown in the Appendix of [1], eq. (4) provides a reasonable estimate also for rectangular or triangular distributions. For verifying eq. (4) we modified the CAP test signal (described in section A.3) such that peak latency was randomly displaced with statistic distribution *δ*_*n*_(*T*). We set *τ* to two test values (i.e. 50 µs and 500 µs, corresponding to the border frequencies *f*_*B*_ 1.6 kHz and 160 Hz, respectively). Since the original CAP template was digitized at 5 µs, we were able to perform accurate time shifts by applying linear interpolation between data points. For studying only the effect of activation jitter, we added no background noise to the CAP signal.

At the low jitter level (*τ* = 50 µs, *f*_*B*_ = 1.6 kHz) evoked activity was accurately captured as reflected by an error of −0.05 dB ± 0.05 dB below 500 Hz. At the high jitter level (*τ* = 500 µs, *f*_*B*_ = 160 Hz) large errors were observed at some 100 Hz (see Figure A.4). The distortion of the signal spectrum by irregular activation generated spectral components, which were reflected as background activity. Below *f*_*B*_ = 160 Hz *N*-FTA captured evoked activity well and background components were well below evoked activity. Above, *f*_*B*_ the error in the evoked activity increased rapidly. At 450 Hz the estimated error in evoked activity was −8 dB and spectral power was almost entirely shifted to background activity.

It must be emphasized that such a high jitter lever of *τ* = 500 µs is far above a realistic value for an experimental setting, since this would induce time shifts of more than 1 ms. This high level was assigned to the synthetic data for illustration purposes. The lower jitter level of *τ* = 50 µs appears being more realistic. In our experimental setting, the width of the stimulation pulse was 200 µs. Assuming a rectangular distribution of this width, we obtain a standard deviation of 58 µs (see Appendix of [1], equation (32)). Thus, our analysis shows that for frequencies below 1 kHz irregular activation or activation jitter may have minor impact on the results, when investigating experimental data.

**Figure A.4:**
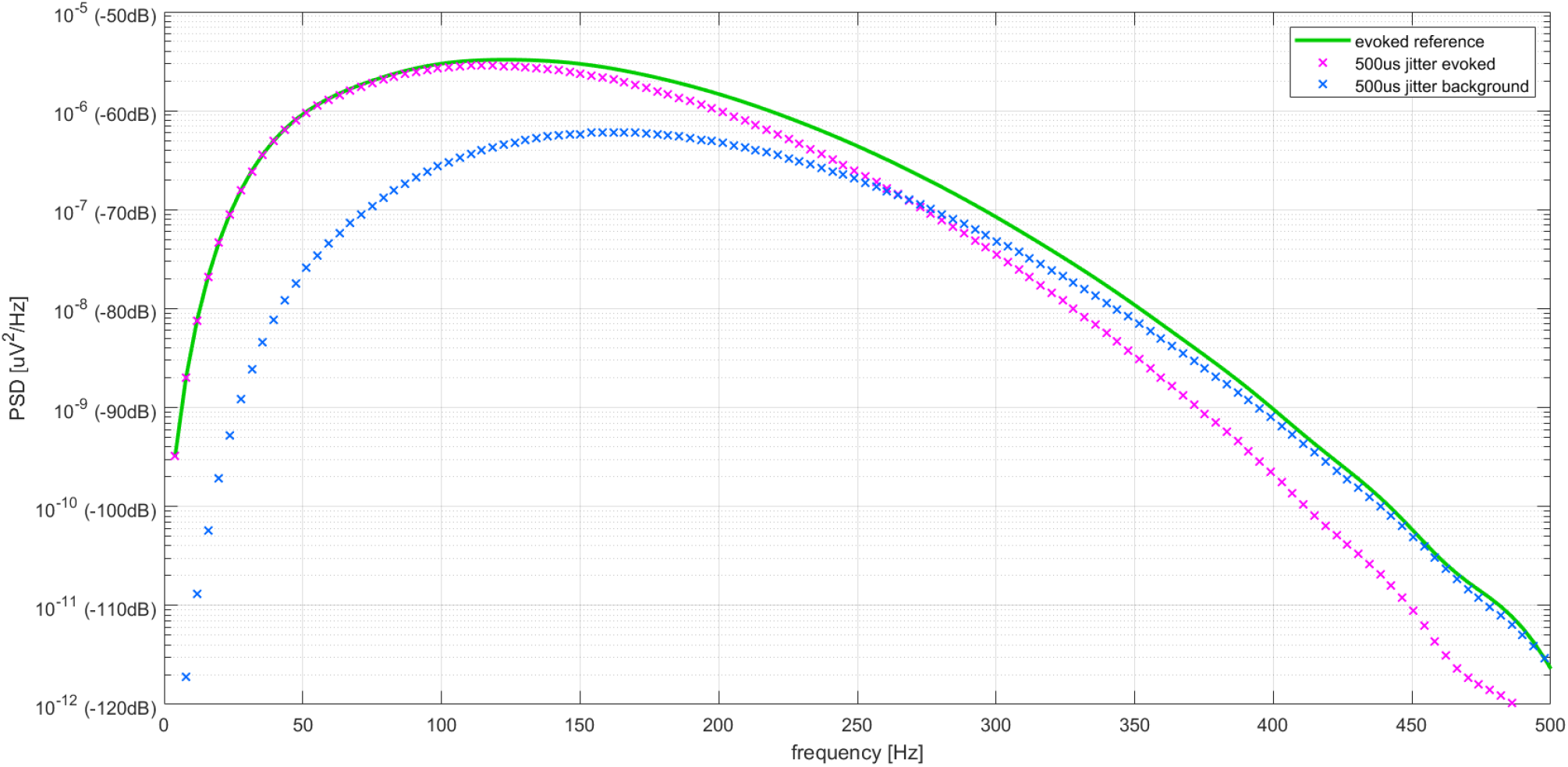
Application of *N*-FTA to a test signal with activation jitter. An unrealistically high jitter level of 500 µs was chosen for illustration (see text).

###### Activation Drift

Physiological effects like, e.g., a drift in body temperature, may cause a drift in the activation timing [2]. As a basic model, a constant drift of activation timing was considered. As it can be seen from Figure A.3, this shifts harmonic peaks. We denote the total time shift occurring over *N* repetitions by Δ*T* and obtain for the change Δ*f*_*E*_ of the evoked frequency

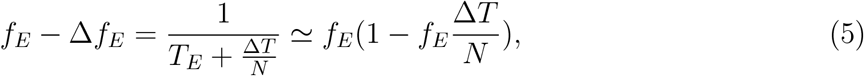

yielding

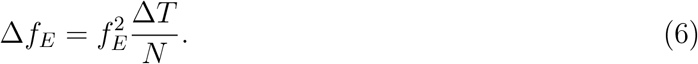

We neglect the negative sign since we are interested in the absolute change in frequency. At each evoked harmonic *k*, the shift in frequency gets multiplied by *k*

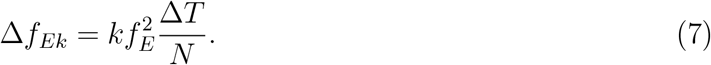

Thus, frequency shifts increase with index *k*. We aim to estimate a maximal allowable change in frequency Δ*f*_*max*_ at the highest frequency of interest *f*_*max*_ = *k*_*max*_*f*_*E*_

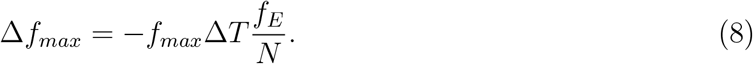

The term 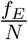 defines the frequency resolution obtained for the finite periodic interval. The maximal change in frequency should not exceed half the frequency resolution

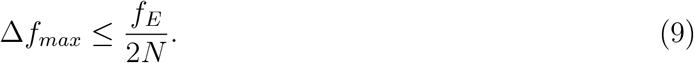

We obtain by inserting (9) into (8)

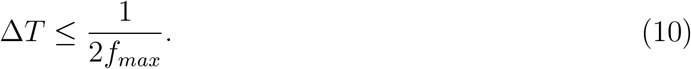

Thus, when aiming to analyze frequency up to 1 kHz, activation drift must be less than 0.5 ms, which is at the upper limit of the clinically accepted reproducibility for latency. We simulated a time shift Δ*T* = 0.5 ms. The errors in the obtained evoked responses were small (−0.19 dB ± 0.17 dB).

###### Variation of Amplitude

The amplitude of individual responses may vary over time. This can be considered as an amplitude modulation. In the frequency domain, amplitude modulation was reflected by side-bands in each spectral harmonic *f*_*En*_ (see Figure A.3).

We simulated such a variation of amplitude by a linear decay from a high initial amplitude to a lower amplitude after *N* repetitions. We chose a factor of 1.5 at the beginning and a factor of 0.5 after *N* cycles, thereby investigating a relatively strong variation at an unchanged mean amplitude. The errors in the obtained evoked responses were −0.01 dB ± 0.01 dB. The irregular activity due to amplitude modulation (side bands) was reflected by a background activity being approximately 10 dB below the evoked level.

###### Summary

Theoretical treatment and numerical simulations showed that inter-cycle variations in repetitions have stronger influence on the high frequency portion of the spectrum, as compared to low frequencies. Results suggest a rather small influence of inter-cycle variations on *N*-FTA below approximately 1 kHz for the chosen experimental setting.

### B Stimulus Cancellation

Stimulation artifacts were short pulses of a few ms duration. Figure B.1 provides a sketch of the cancellation approach. We observed from the experimental data that the artifact can be split in two phases. An initial phase A of high amplitude and significant high frequency content. Since the stimulation pulse had a duration of 200 µs it contained frequency content beyond the 4.8 kHz Nyquist frequency of AD-conversion. Thus, within phase A the stimulus artifact was under-sampled. In a later phase B the artifact displayed a relatively smooth asymptotic return to zero which was accurately captured.

The timing of the stimulation pulses was obtained from a trigger channel. At 9.6 kHz sample rate each 200 µs stimulation pulse contained two non-zero samples. The index of the first non-zero sample defined time zero for the individual stimulus (index *m*_*z*_). Based on the observations from the experimental data, we selected a width of *M*_*A*_ = 10 samples for interval A (window from *m*_*z*_ to *m*_*z*_ + *M*_*A*_, i.e., approximately 1.1 ms). For interval B we selected a width of *M*_*B*_ = 45 (window from *m*_*z*_ + *M*_*A*_ + 1 to *m*_*z*_ + *M*_*A*_ + *M*_*B*_, i.e., approximately 4.7 ms).

**Figure B.1:**
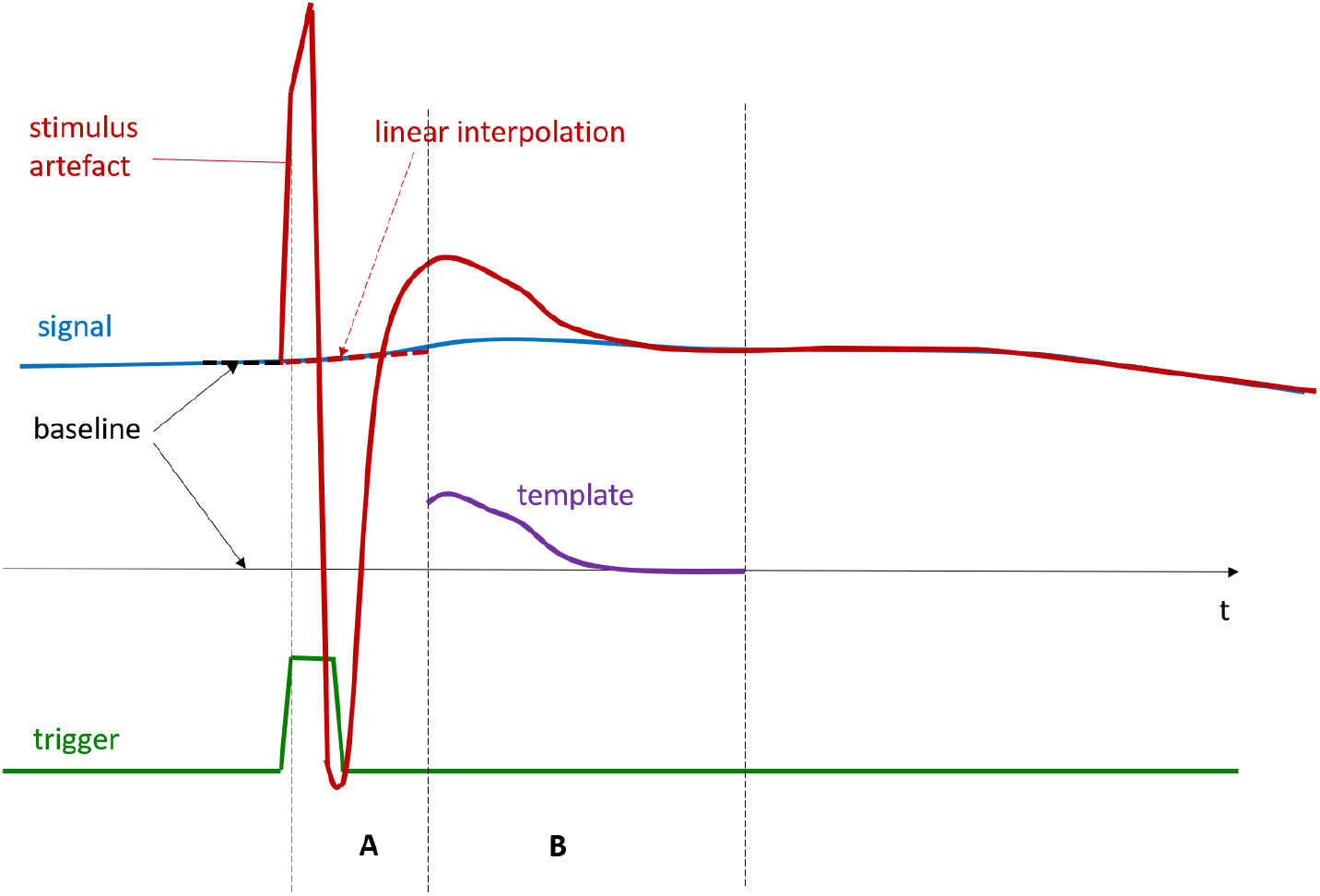
Illustration of the approach applied for stimulus cancellation. In the raw data a stimulus artifact (red) was superimposed on the signal (blue). Timing of the stimulation pulses was obtained from a trigger channel (green). The stimulation pulse was split into an early phase A and a later phase B. For phase B a stimulus template was subtracted from the signal. In phase A linear interpolation was performed (see text for details).

We computed a stimulus template within interval B by averaging the signal of all *N* intervals B. We estimated the baseline of this artifact by averaging five stimuli ahead of all *N* stimuli. We subtracted the baseline corrected template in each interval B from the recorded data. We “bridged” the signals in intervals A by performing linear interpolation. Figure B.2 shows example data. Interstimulus variations of high frequency can be observed within intervals A. For the depicted example the stimulus amplitude near the left border of interval B was approximately 7 µV. This appears as a small visible distance between raw data and preprocessed data in the figure. However, due to the small amplitudes of SEPs it was important to accurately remove such small stimulus components before performing spectral analysis.

### C Bandwidth Estimates

#### C.1 MATLAB Function filtfilt.m

The MATLAB function `filtfilt.m` provides a zero-phase filter by applying an infinite impulse response (IIR) filter twice. The IIR filter is applied once in a forward time direction and once with reversed time direction. Thus, phase shifts compensate each other providing a zero-phase filter.

When using `filtfilt.m` in a first step the IIR coefficients are computed which correspond to a selected −3 dB bandwidth for single application of the filer (i.e., MATLAB function `filter.m`). Due to the double application of the filter the originally selected bandwidth becomes a −6 dB bandwidth by using `filtfilt.m`. Furthermore, the filter order is doubled by using `filtfilt.m`. In this section, we provide also the `filtfilt.m` −3 dB bandwidth, for allowing a direct comparison with standard filter types.

##### High-Pass Filter

For the MF-band setting of short latency SEPs we used a 2^nd^-order Butterworth filter. IIR coefficients were computed for an original −3 dB corner frequency of 14.4 Hz. We computed the frequency response for single (function `filter.m`) and double (`filtfilt.m`) application of these coefficients (see Figure C.1). We obtained a −3 dB corner frequency of 17.99 Hz when using `filtfilt.m`.

##### HFO Band-Pass Filter

For investigating high frequency oscillations (HFOs) the identical zero-phase band-pass as described in [3] was implemented. Here, the IIR coefficients for a 3^rd^ order Butterworth 400 Hz to 800 Hz band-pass filter were used. We computed the frequency response for single (function `filter.m`) and double (`filtfilt.m`) application of these coefficients (see Figure C.1). We obtained −3 dB corner frequencies of 419 Hz and 765 Hz when using `filtfilt.m`.

#### C.2 HFO Bandwidth & PSD-Level

The aim of this section is to obtain coarse estimates of expected power level of near 600 Hz HFOs. HFOs *φ*_*M*_(*t*) may be coarsely approximated by a Morlet wavelet of amplitude *V*_*M*_, frequency *f*_*M*_ and time constant *τ*_*M*_ [6]

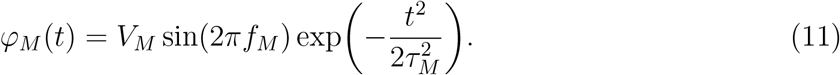

Figure C.2 displays the test signal which was obtained by choosing the parameters *V*_*M*_ = 0.25 µV, *f*_*M*_ = 600 Hz and *τ* = 3 ms. We sampled these data at *f*_*S*_ = 9.6 kHz within an interval of length *T*_*E*_ (*f*_*E*_ = 3.95 Hz). We numerically computed its PSD representation as also shown in the figure.

We aim for obtaining a “rule of thumb” estimate for the PSD level near the spectral peak, by using rectangular windows. The average power of the Morlet wavelet 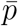 in one evoked cycle can be estimated from a rectangular window of amplitude *V*_*M*_ and width 2*τ*_*M*_ by

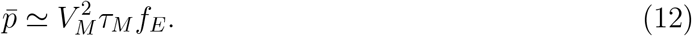

A coarse estimation of the bandwidth is obtained using equation (30) from the Appendix of [1]. For a rectangular pulse, the first zero crossing in the Fourier transform occurs at the inverse of the width of the window, i.e., at 2*τ*_*M*_. The effective bandwidth is always somewhat smaller than the interval defined by the zero crossing. As an estimate, we use the inverse of three times *τ*_*M*_

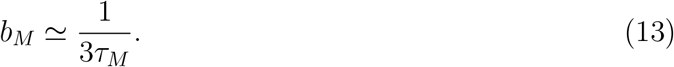

The PSD level of the HFO burst is the ratio of power 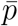 and bandwidth *b*_*M*_. We obtain

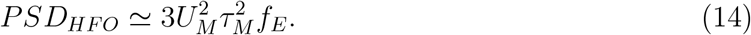

From our own data and the data in [3, 6] we estimated, that *V*_*m*_ may be selected from the range of 0.15 µV to 0.30 µV and *τ*_*m*_ from 2.0 ms to 4.0 ms. From these values we estimated PSD level by −60 dB to −48 dB (relative to the 1 µV^2^/Hz level) for the range of the expected HFOs.

---

**Figure B.2:**
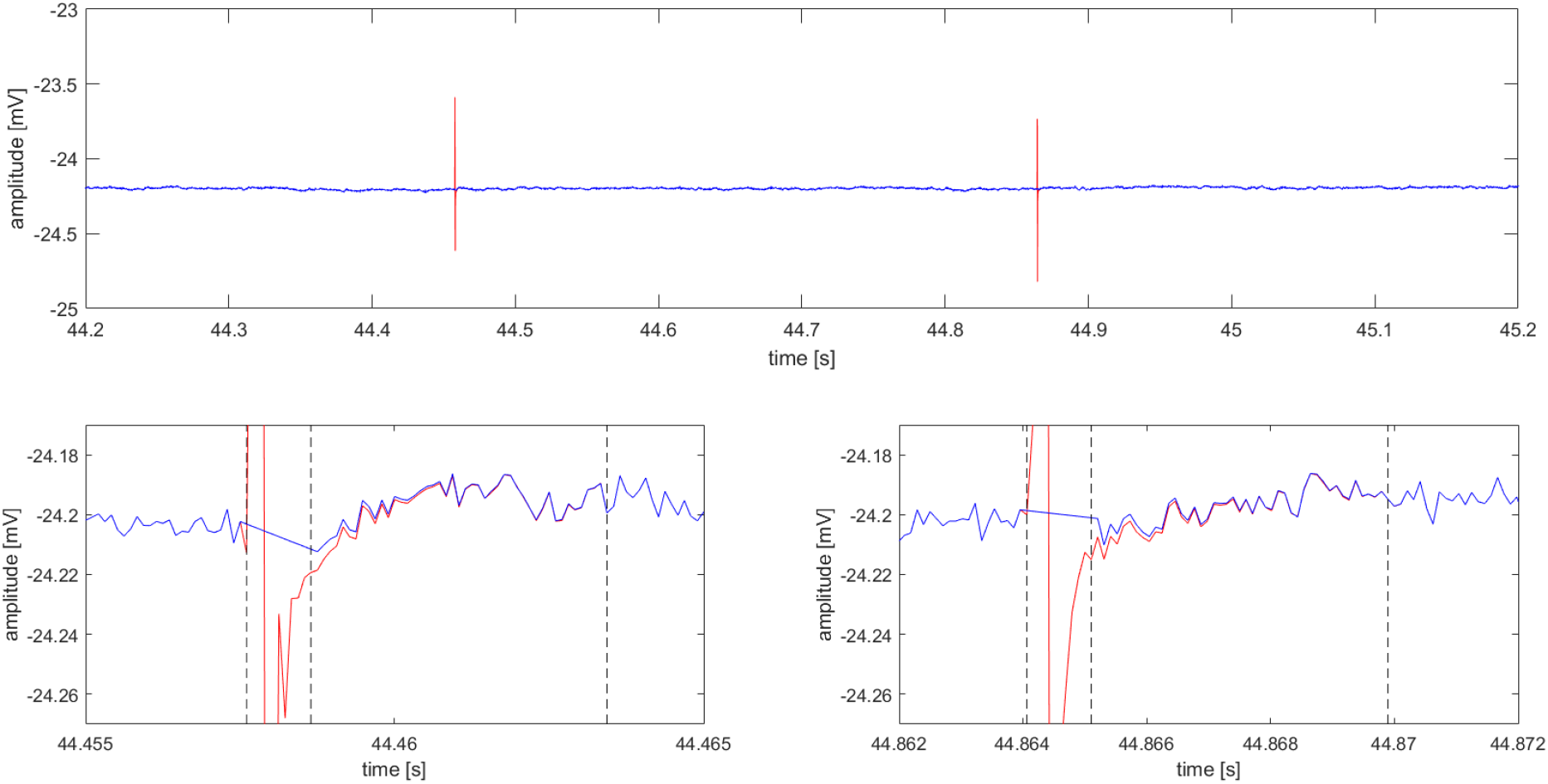
*Top:* Example raw data (red) and signal after stimulus cancellation (blue). A time interval containing two stimuli was shown. True DC recording of the signal displayed a remarkable DC offset (approximately −24 mV). The peak-to-peak amplitude of the artifacts was approximately −1 mV. *Bottom left:* The first stimulation artifact from the top panel was magnified. Vertical dashed lines correspond to time windows A and B in Figure B.1. *Bottom right:* The second stimulation artifact from the top panel was magnified and shown in the same style.

**Figure C.1:**
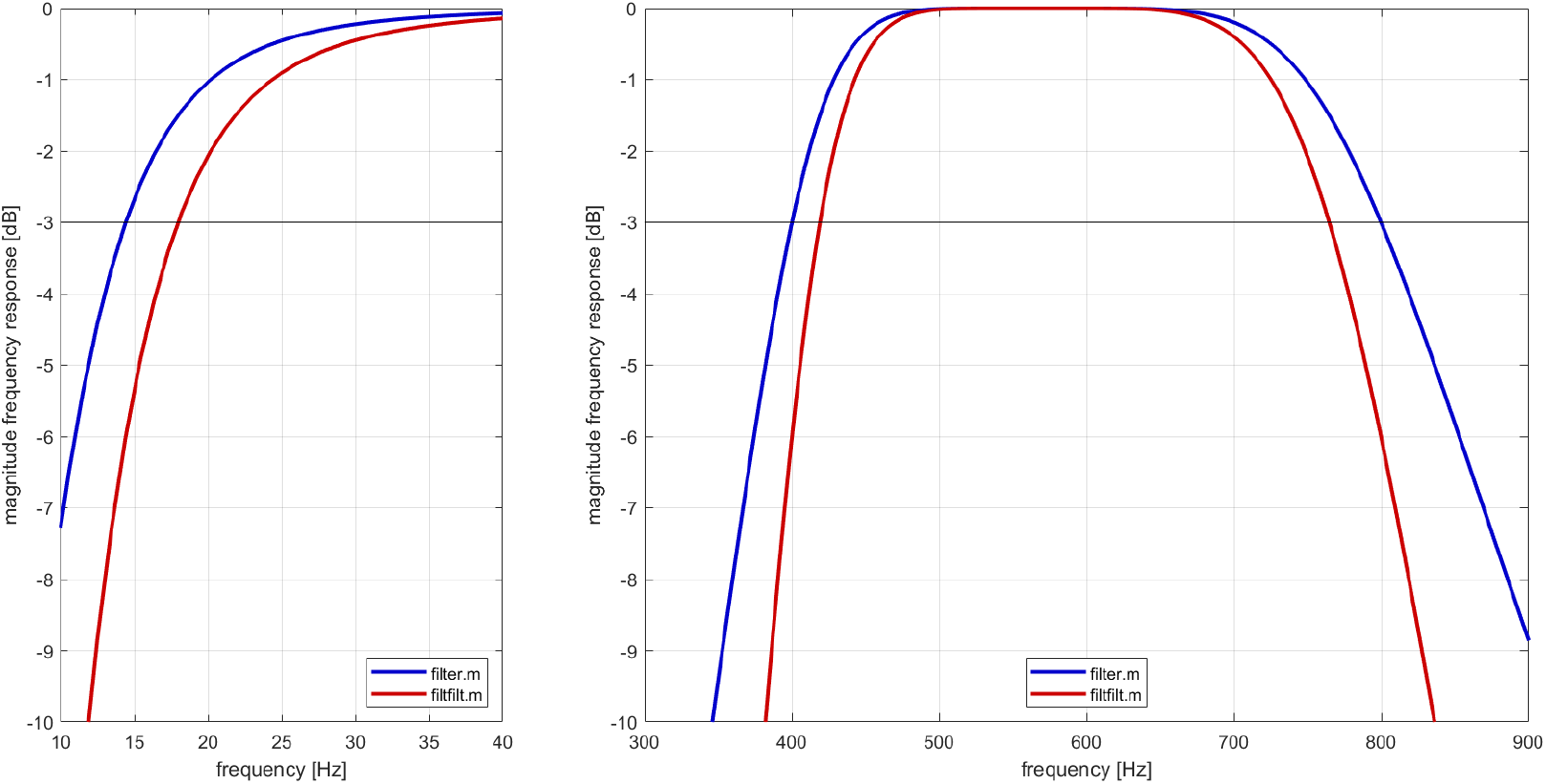
Magnitude of frequency responses obtained by using functions MATLAB `filter.m` and `filtfilt.m` with the same IIR-coefficients. The −3 dB level is indicated by a narrow line. *Left:* Low-pass filter for short latency SEPs. *Right:* Band-pass for HFOs.

**Figure C.2:**
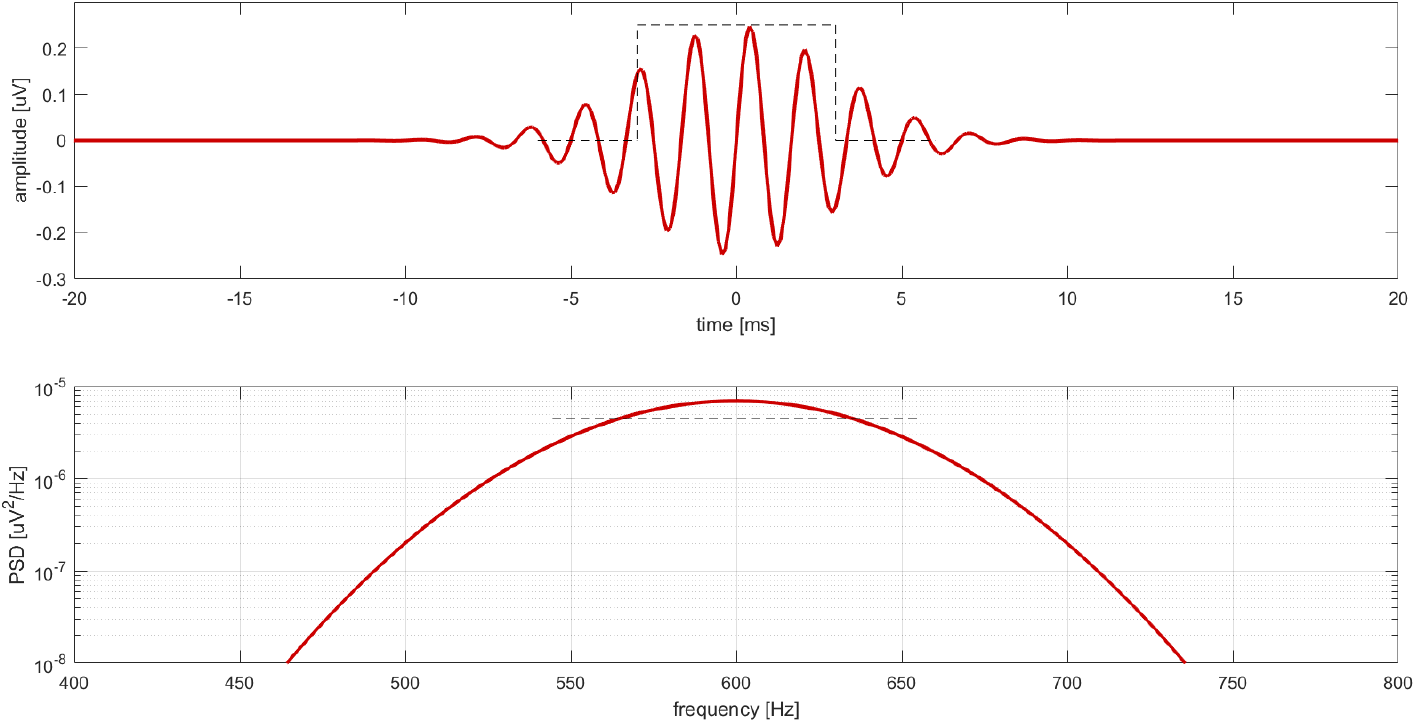
*Top:* Approximation of an HFO by a Morlet wavelet with the parameters listed in the text (red trace). The dashed lines indicate the rectangular window of equal power. *Bottom:* PSD of the Morlet HFO (red trace). The dashed line marks the PSD level estimated from the rectangular window.

## Notes

### Competing Interest Statement

The authors have declared no competing interest.

